# Atomistic simulations reveal sub-*µ*s contact dynamics in MUT-16 condensates

**DOI:** 10.64898/2026.03.30.715404

**Authors:** Kumar Gaurav, Lucia Baltz, Diego Javier Páez-Moscoso, René F. Ketting, Lukas S. Stelzl

**Affiliations:** Institute of Molecular Physiology, Johannes Gutenberg University Mainz, Germany; Institute of Molecular Biology (IMB), Mainz; Institute of Quantitative and Computational Biosciences (IQCB), Johannes Gutenberg University Mainz, Germany; Max Planck Graduate Center (MPGC) Mainz, Germany

**Author notes:** These authors contributed equally to this work (Kumar Gaurav, Lucia Baltz).

## Abstract

Phase separation of proteins gives rise to biomolecular condensates, which function as membraneless organelles that spatially and temporally organize cellular functions. Such condensates are often formed by intrinsically disordered regions of proteins (IDRs), whose multivalent and transient interactions govern condensate structure and dynamics. However, elucidating the molecular determinants of these interactions at atomistic resolution remains challenging. Here, we present a total of 10 *µ*s of atomistic molecular dynamics simulations of a phase-separated condensate formed by the foci-forming region (FFR) of MUT-16. MUT-16 serves as a scaffold of the *Mutator foci* germ granules in *Caenorhabditis elegans* and is essential for transposon silencing. MUT-16 FFR is enriched in polar uncharged (Gln, Asn), charged, aromatic, and Pro residues, raising the question of how these amino acids interact within condensates. We find that most contacts are short lived, typically breaking within a few nanoseconds (ns), with a median life time of 9.8 ns. A smaller fraction persist for much longer timescales (*>* 100 ns). We characterized the relative contributions of different amino acids and specific interaction types, including hydrogen bonding, cation–*π* interactions, *π*–*π* stacking, and salt bridges and theirs dynamics. We further examined the roles of water and ions in modulating condensate interactions, including ion-mediated bridging between similarly charged residues. Our results reveal that salt bridges, cation-*π* interactions, Na^+^ ions, and water in the condensate are key determinants of contact dynamics in MUT-16 FFR condensates. In parallel, we show that these condensates exhibit upper-critical solution temperature (UCST) phase behavior *in vitro*, providing a coherent framework to explain both the loss of *Mutator foci* at elevated temperatures *in vivo* and the scaffolding role of MUT-16 at lower temperatures.

## Introduction

The formation of phase separated biomolecular condensates has emerged as a fundamental organizing principle in cells.^1–3^ These condensates give rise to membraneless organelles, including the nucleolus and stress granules.^4–6^ In germ cells, multiple types of biomolecular condensates, collectively termed germ granules, have been described. *Mutator foci* are a nematode-specific germ granule type membraneless organelle involved in small RNA biogenesis for genome surveillance, with MUT-16 as the primary scaffold protein.^7^ A key molecular feature underlying condensate formation is the prevalence of intrinsically disordered proteins (IDPs) or proteins containing intrinsically disordered regions (IDRs).^8,9^ Unlike folded proteins, IDPs lack stable secondary and tertiary structure, allowing them to sample highly dynamic conformational ensembles^10^ and engage in transient and multivalent interactions that underpin their phase separation.^11^ Consequently, biomolecular condensates formed by IDPs are central to a wide range of biological functions,^12–14^ while their dysregulation has been implicated in pathological conditions such as neurodegenerative diseases.^15^ Beyond their biological relevance, the physicochemical principles governing IDP-driven condensates provide a foundation for the rational design of biomimetic and stimuli-responsive materials,^16^ underscoring the importance of understanding condensate formation and dynamics at the molecular level.

Despite significant advances in experimental techniques for probing biomolecular condensates, directly resolving their internal dynamics, interaction networks, and the specific molecular interactions driving phase separation remains challenging.^8,17–20^ Accumulating evidence suggests that protein phase separation is governed by a complex interplay of noncovalent interactions, including electrostatic, hydrophobic, *π*–*π*, and cation–*π* interactions between amino acid residues.^19,21–30^ The physical chemistry of these non-covalent interactions has been well characterized in model systems,^31–35^ but their roles in biomolecular condensates remain incompletely understood. In addition to direct protein–protein interactions, ion-mediated interactions and water-mediated effects have been shown to play essential roles in modulating condensate stability and dynamics.^19,29,36^ Atomistic molecular dynamics simulations with explicit solvent provide a powerful framework for dissecting these interactions at molecular resolution, offering insights into the chemical specificity and dynamics that are difficult to access experimentally.^19,29,37–40^ Molecular dynamics simulation can resolve directly how different amino acids interact^20,30,38,41^ and the dynamics of their interactions can be quantified.^30,42,43^ Previous molecular dynamics studies have shown that biomolecular condensates are characterized by heterogeneous networks^22,26^ of both short- and long-lived intermolecular interactions. In a pioneering atomistic investigation, Zheng *et al.* revealed the molecular interactions that underpin protein phase separation.^19^ More broadly, Rekhi *et al.* combined atomistic simulations with experiments to show that protein phase separation can be supported by a diverse repertoire of interactions extending beyond, e.g., aromatic *π*-mediated contacts involving Arg.^38^ Rekhi *et al.* also reported highly heterogeneous contact dynamics, with some contacts breaking and forming repeatedly. Galvanetto *et al.* reported tracking the dynamics of contacts in simulations of a coacervate formed by negatively charged prothymosin *α* and positively charged histone H1, highlighting the central role of salt bridges and ion-mediated interactions^20^ and reported that contacts typically break on the timescale of ns, while a small fraction of contacts persists for longer times. Recently, Galvanetto *et al.* have identified dynamic reshuffling of such non-covalent interactions in multiple condensates and linked this to the mesoscopic properties of condensates.^43^ Along similar lines, our previous work on the MUT-16 system identified long-lived Tyr–Arg contacts stabilized by cation–*π* interactions within a dense cluster of chains of a disordered segment of MUT-16, although this region alone does not undergo phase separation.^39^ Despite these advances,^19,20,38,43^ the relative lifetimes of residue-level interactions and the role of ion and water-mediated bridging in shaping the dynamics in condensate remain incompletely understood.

However, the large size of condensate systems and long simulation durations required to achieve equilibrium coexistence of dilute and dense phases pose a major challenge for fully atomistic simulations. This limitation can be addressed by employing coarse-grained simulations to generate equilibrated, phase-separated configurations,^19,44–46^ which can then be backmapped to all-atom representations.^19,47,48^ This multiscale strategy enables detailed interrogation of atomic-level interaction patterns while preserving access to larger scale condensate organization.^19,38^ In practice, however, such comprehensive analyses impose substantial computational overhead, as individual interaction metrics are often evaluated using distinct analysis tools that require separate passes over long simulation trajectories.^41,49^ This fragmentation not only increases computational cost but also complicates data provenance, reproducibility, and reuse, making adherence to FAIR data principles^50^ challenging in large-scale molecular simulation studies.

MUT-16 is a scaffolding protein that nucleates the formation of a membraneless organelle known as the *Mutator foci* in *Caenorhabditis elegans*. These condensates plays a central role in the amplification of small interfering RNAs (siRNAs), a critical step in RNA silencing.^7,51^ RNA silencing, commonly known as RNA interference (RNAi), is a conserved gene-regulation mechanism that safeguards genome integrity by suppressing viruses and transposable elements and is essential for processes including gametogenesis, chromosome segregation, and development.^52,53^ The ability of MUT-16 to assemble into *Mutator foci* is therefore fundamental to the proper execution of RNAi-mediated gene regulation. *Mutator foci* dissolve as the ambient temperature is increased, which has been interpreted as upper critical solution temperature (UCST) behavior of protein condensates. MUT-16 contains a large intrinsically disordered region (IDR), within which a segment spanning residues 773-945 has been identified as the foci-forming region (FFR) that is necessary for MUT-16 phase separation.^51^ Deletion of this region abolishes *Mutator foci* formation *in vivo*, while *in vitro* experiments demonstrate that the FFR alone is sufficient to undergo liquid–liquid phase separation and form condensates.^39^ Notably, an adjacent region of MUT-16 is required for recruiting RNA-processing machinery^54^ but is dispensable for foci formation; this region does not phase separate *in vitro*, a distinction that is recapitulated in residue-resolved coarse-grained and Martini3 simulations.^39^ Despite these advances, the atomistic interactions and molecular mechanisms that drive phase separation of the MUT-16 FFR remain poorly understood.

In this study, we use atomistic simulations to resolve the dynamic interactions within the phase-separated MUT-16 FFR condensate. We established that MUT-16 condensates are (1) a representative model system for protein condensates by sequence analysis and (2) for *Muta-tor foci in vivo* by characterizing the temperature-dependence of MUT-16 phase separation. We then elucidated the molecular interactions governing phase separation of the MUT-16 FFR at atomistic resolution (Movie S1) by performing a comprehensive, residue-resolved analysis of contact frequencies and relating these interaction propensities to the corresponding lifetimes of residue pairs. In addition, we quantified the relative contributions of specific non-covalent interactions, including hydrogen bonding, salt bridges, cation–*π* interactions, *π*–*π* stacking, and determined how their fraction of contacts and lifetimes shape condensate dynamics. Beyond protein–protein interactions, we examined the association of Na^+^ and Cl^−^ions with amino acid side chains and backbones, analyzed the persistence of ion–residue interactions, and investigated the role of counter-ions in mediating interactions between similarly charged residues. Finally, we characterize the distribution and dynamics of water within the condensate, comparing hydration in the condensate interior and at the interface, and evaluate how water molecules bridge interactions among acidic, basic, and polar residues. To mitigate the computational overhead associated with large-scale atomistic analyses, we employed parallelization on high-performance computing (HPC) infrastructure also for the residue-resolved contact analysis, implementing a multi-step downstream analysis pipeline that operates on a unified contact record (https://github.com/luhtzia/cascade_computing).

## Methods

### Sequence analysis of MUT-16 FFR sequence

To contextualize the MUT-16 FFR within the broader landscape of IDPs, we analyzed its sequence similarity and amino-acid composition relative to a comprehensive dataset of 28,058 human intrinsically disordered regions (IDRs) compiled by Tesei *et al.*^10^ For comparison, the same analyses were performed for the low-complexity domain (LCD) of the human FUS protein, a well-established model system for protein phase separation.^19,55^ Embedding-based alignment (EBA)^56^ was used to quantify sequence-level similarity between MUT-16 FFR and human IDRs. Protein sequence embeddings were generated using the ESM-2 protein language model (facebook/esm2_t12_35M_UR50D).^57^ For each human IDR, an embedding-based similarity score (EBA_min_) was computed with respect to MUT-16 FFR. An identical procedure was applied to compute EBA_min_ scores for the FUS LCD against the same human IDR dataset. All EBA calculations were performed following the reference implementation provided at https://git.scicore.unibas.ch/schwede/EBA. To assess compositional similarity independent of sequence order, we additionally computed the root-mean-square error (RMSE) between amino-acid composition vectors. For each sequence, the abundance of the 20 standard amino acids was represented as a composition vector. RMSE values were then calculated between the composition vectors of each human IDR and those of MUT-16 FFR or FUS LCD. This metric provides a quantitative measure of similarity in residue composition and enables direct comparison of MUT-16 FFR and FUS LCD with the broader human IDR proteome.

### Temperature-dependent phase separation experiments

#### Protein labeling with Atto488-maleimide

Recombinant MBP-3C-MUT-16-6xHis 773–944 aa (FFR) protein was purified as described in.^39^ The protein was then buffer-exchanged into labeling buffer (100 mM NaPi, pH 7.1, 150 mM NaCl) using Zeba Spin Desalting Columns (Thermo Scientific, A57760) following the manufacturer’s instructions. The cysteines were reduced by the addition of 2 mM TCEP and incubated on ice for 10 minutes. Atto488-maleimide (ATTO-TEC, AD488-41) was then added in a 2.5-fold molar excess relative to the protein and incubated at 4°C overnight. To quench the reaction, 4 mM DTT was added, and the free dye was removed twice using Zeba Spin Desalting Columns (Thermo Scientific, A57760) following the manufacturer’s instructions.

#### Preparation of emulsions

The concentration of the purified MUT-16 construct MUT-16 773–944 aa (FFR) was determined via extinction coefficient measurements at 280 nm using a DeNovix DS-11 spec-trophotometer (Biozym) and diluted to the desired concentration in storage buffer (20 mM Tris/HCl, pH 7.5, 150 mM NaCl, 10% (v/v) glycerol, and 2 mM DTT). Atto488-labeled MUT-16 recombinant protein was also added, representing 1% of the total protein in the reaction, to better delineate the contour of the condensate during analysis. Finally, conden-sate formation was induced by cleaving the MBP tag using 3C protease (1 mg/mL, in-house produced) at a 1:100 (w/w) ratio of 3C protease to the corresponding MUT-16 fragment. Immediately afterward, PicoSurf oil (2% (w/w) in Novec 7500, Sphere Fluidics, UK) was added at 2x the solution volume to encapsulate the reaction to have a define volume of the reaction. Each buffer-oil mixture was agitated using a 10 µL pipette and 10 µL tips until the desired size distribution of water-in-oil droplets was achieved. The reactions were performed in 8-strip PCR tubes (Multiply-µstrip Pro 8-strip, Sarstedt, REF 72.991.002). The specific titration series of MUT-16 773–944 aa (FFR)[20 µM, 15 µM, 10 µM], shown in Fig. 3B, were then mounted on a temperature-controlled microscope stage.

**Figure 1:**
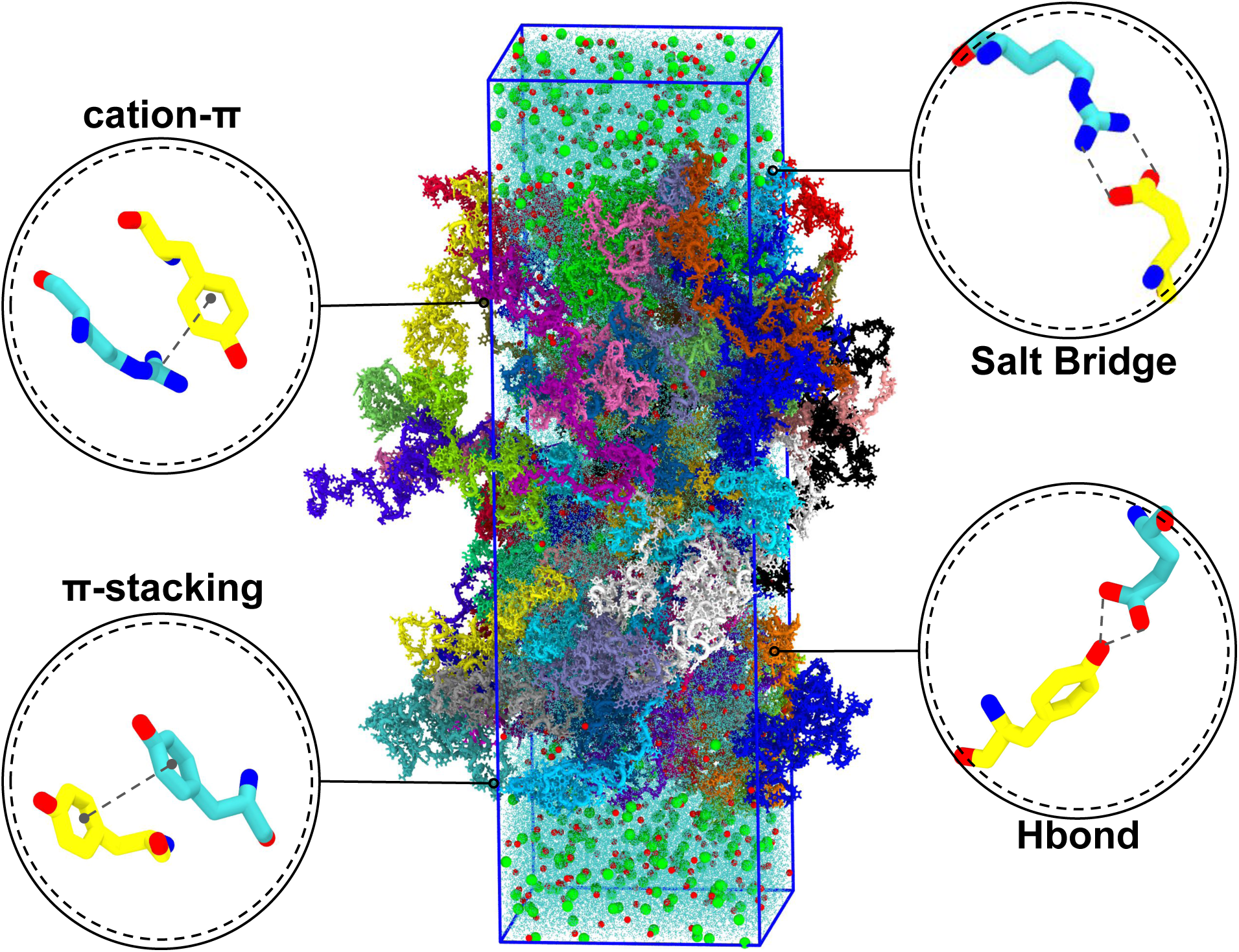
Simulation slab box containing MUT-16 FFR chains, each represented by a distinct color. The protein chains are solvated in water (cyan) with ions, Na^+^ (red) and Cl^−^ (green). Representative specific interactions observed in the simulation are highlighted, including cation-*π* interactions between the guanidinium group of Arg and the phenol group of Tyr, *π*-*π* stacking between phenol groups of Tyr residues, salt bridges between the guanidinium group of Arg and the side-chain carboxylate group of Glu, and hydrogen bonds between the hydroxyl group of Tyr and the carboxylate group of Glu.

**Figure 2:**
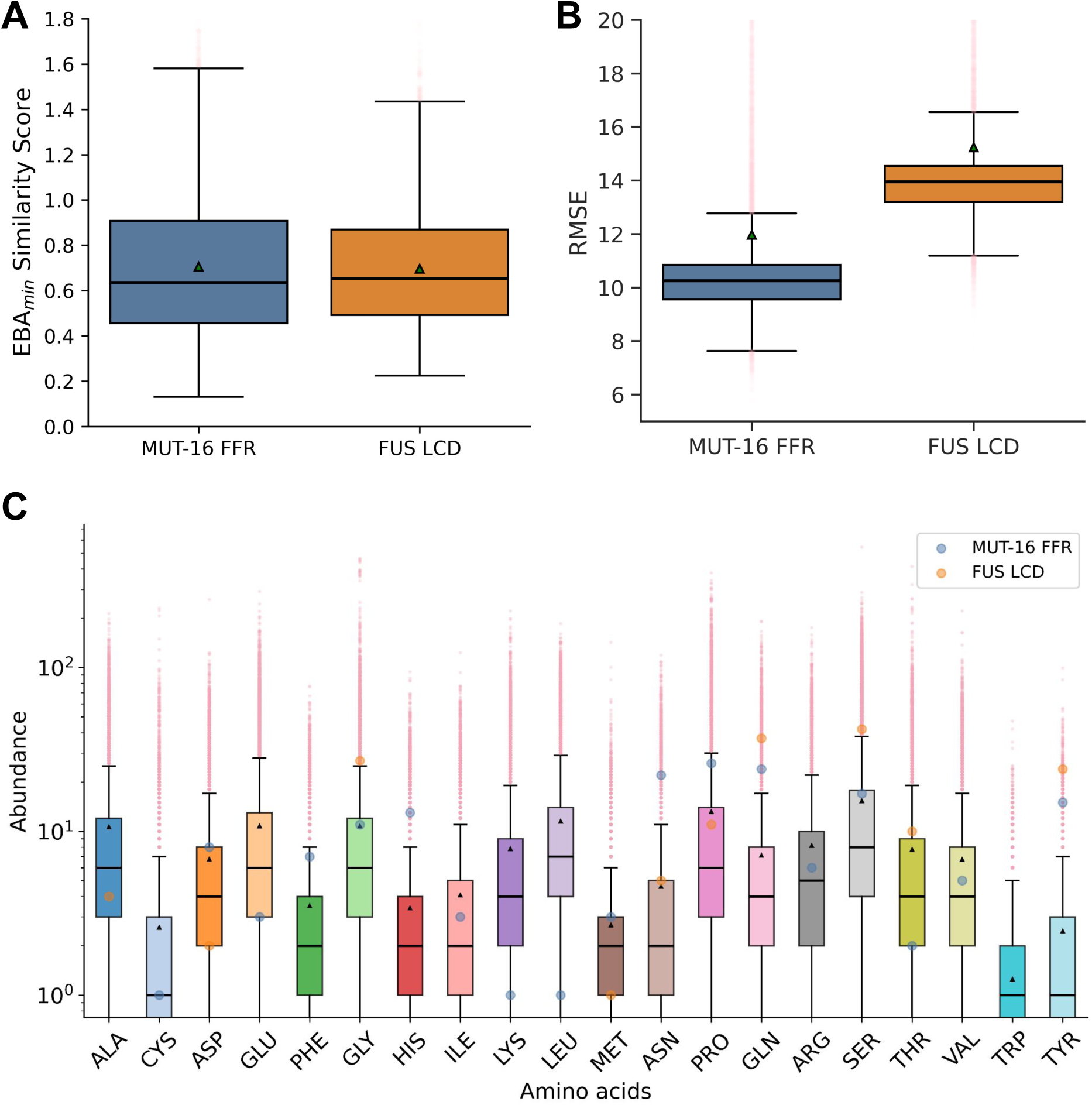
Sequence and compositional similarity of human IDRs to MUT-16 FFR and FUS LCD. **A.** Box-and-whisker plots showing the distribution of EBA_min_ similarity scores for human IDRs, evaluated with respect to the MUT-16 FFR (blue) and the FUS LCD (orange). The median is indicated by the horizontal line and the mean by a green triangle; outliers are shown as translucent pink stars. **B.** Box-and-whisker plots showing the distribution of RMSE values quantifying compositional similarity of human IDRs relative to the MUT-16 FFR (blue) and the FUS LCD (orange). **C.** Box-and-whisker plots showing residue abundance distributions across the same set of human IDRs, with residue abundances for the MUT-16 FFR (blue) and the FUS LCD (orange) overlaid.

**Figure 3:**
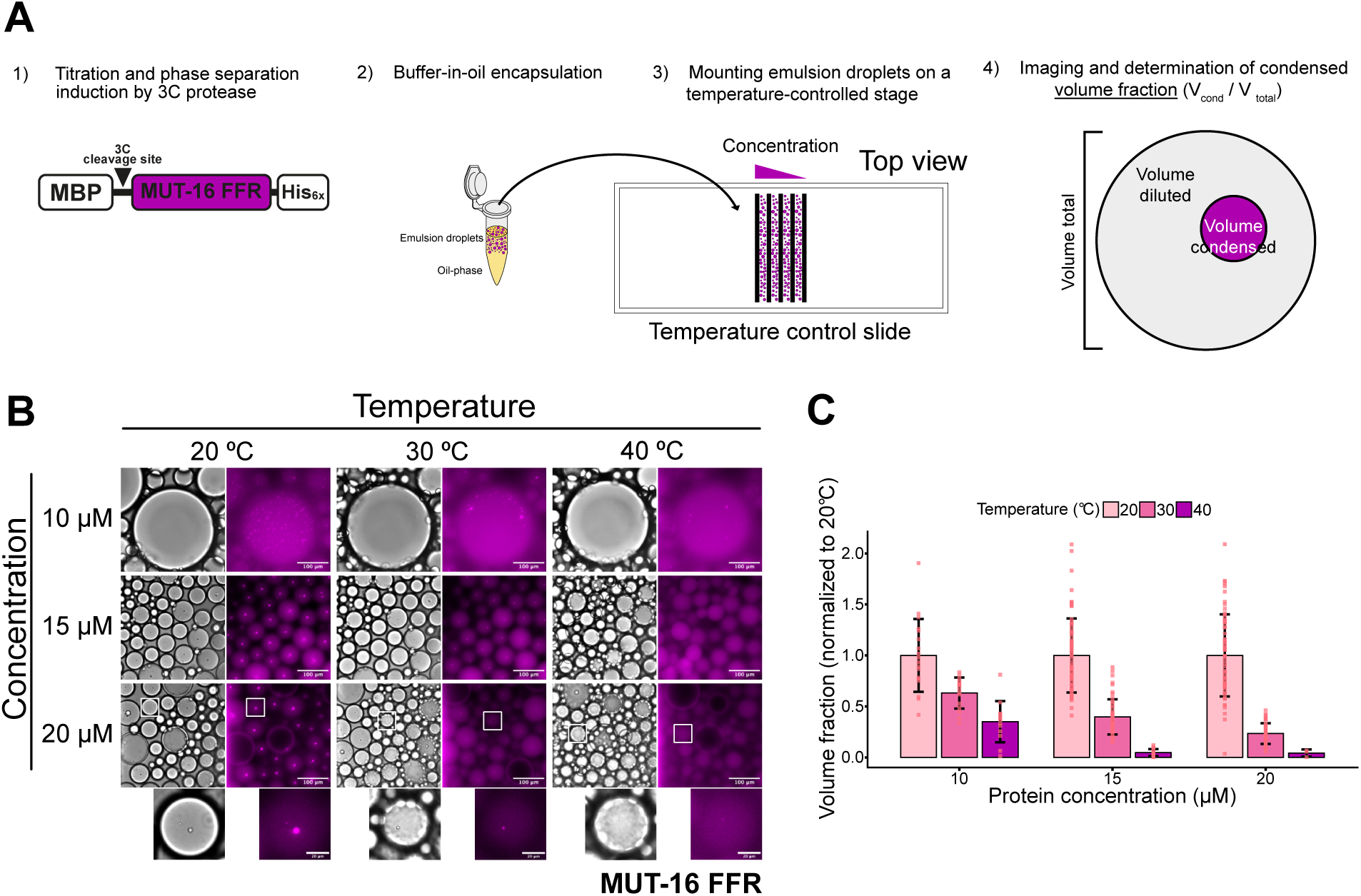
MUT-16 FFR is a low-temperature condensation (UCST) protein. **(A)** Schematic representation of the experimental workflow for the *inPhase* method. In the first step, serial dilutions of MUT-16 fragments are prepared, and phase separation is induced by cleaving the MBP tag using 3C protease. In the second step, the reaction mixture is immediately encapsulated into oil droplets. In the third step, the emulsion droplets are loaded onto a temperature-controlled stage between two Parafilm strips to spatially compartmentalize each condition (e.g., different protein concentrations). In the final step, the droplets are imaged, and the volume fraction (*V*_con_*/V*_tot_) is determined from the condensate and droplet volumes. **(B)** Representative single-plane images of emulsion droplets containing the MUT-16 773–944 aa (FFR) fragment at different concentrations (10 *µ*M, 15 *µ*M, 20 *µ*M) and temperatures (20^◦^C, 30^◦^C, 40^◦^C). Left panels show bright-field images, and right panels show fluorescence images. Scale bar: 100 *µ*m. White boxes in the 20 *µ*M panel indicate regions shown in the zoomed-in images below (scale bar: 20 *µ*m). **(C)** Quantification of temperature effects on condensates. The volume fraction was normalized to the value at 20^◦^C for each protein concentration. Each experiment comprises data points from multiple emulsion droplets. Bars represent mean ± SD (*n* = 47 for 10 *µ*M, *n* = 138 for 15 *µ*M, and *n* = 116 for 20 *µ*M)

#### Sample loading and imaging procedure for emulsions

In order to rapidly and accurately change the temperature on the condensates, a custom-made temperature-controlled microscope stage named Vulcan developed by Anatol W. Fritsch, Juan M. Iglesias-Artola, and Anthony A. Hyman (2024)^58^ was used. Parafilm (Bernis, USA) lanes were created and sandwiched between a sapphire slide and a standard 20x20 mm cover-slip (Roth Karlsruhe) to enable parallel imaging of the titration reaction series. The emulsion solutions were loaded into individual lanes by pipetting, and each lane was sealed to min-imize evaporation and movement of the water-in-oil emulsion. The temperature-controlled microscope stage was set to 25°C for 30 minutes to complete the 3C cleaving reaction and then decreased to 20°C for an additional 30 minutes. Samples were then imaged using a Thunder (Leica) inverted widefield microscope in bright-field and fluorescence modes with a 40x/0.95 air objective. The temperature was then increased in 10°C steps interval until reaching 40°C, with a 30-minute equilibration period before imaging at each interval.

#### Image Analysis and Calculation of volume fraction

The effects in temperature on condensate formation was measured as volume fraction which is the ratio between the volume of condensate(s) over the total volume of the oil droplet. Images were processed using Fiji/ImageJ (2.14.0/1.54f). Bright-field images of each lane were used to detect the contours of the water-in-oil emulsion droplets manually, while fluorescence images were used to identify the contours of the condensates. The analysis assumed an ellipsoid shape for the emulsions, which was measured manually to determine the circumference of each emulsion droplet. The volume of the condensate(s) within each emulsion droplet was determined via image registration, and intensity of the dilute and condensed phases. In brief, after basic image filtering, a Gaussian blur was applied to the image, and a mask of the condensate was generated. This mask was used to estimate the outline of the condensates and, assuming sphericity, to calculate their volumes, which were also manually inspected for accuracy. Then the volume fraction was calculated and normalize to the 20°C for each concentration of recombinant protein.

### Coarse-grained simulations of MUT-16 FFR phase separation

Coarse-grained molecular dynamics simulations of MUT-16 FFR were performed with GRO-MACS (version 2018)^59^ using the Martini3-IDP force field.^44,60^ The MUT-16 FFR (residues 773–944; 172 amino acids)^39^ was initialized from the AlphaFold-predicted^61^ structure (AF-O62011-F1) and converted to a coarse-grained model using the martinize2 pipeline.^62^ A total of 65 copies of the coarse-grained MUT-16 FFR were embedded in a slab-shaped simulation box with dimensions of 12 × 12 × 60 nm^3^ and solvated using the insane.py script.^63^ System neutrality was ensured, and additional Na^+^ and Cl^−^ ions were added to reach a physiological salt concentration of 0.15 M. Periodic boundary conditions were applied in all dimensions. The system was first subjected to energy minimization using the steepest-descent algorithm, followed by a multi-stage equilibration protocol. Initially, solvent and ions were equilibrated in the NVT ensemble while restraining the protein coordinates. These re-straints were subsequently released, and further equilibration was carried out in the NVT ensemble to allow protein relaxation. The systems were then equilibrated under NPT con-ditions using a 20 fs time step. Production simulations were finally conducted in the NPT ensemble. Throughout the simulations, the temperature was maintained at 300 K using the Bussi–Donadio–Parrinello velocity-rescaling thermostat,^64^ and pressure was maintained at 1 bar with the Parrinello–Rahman barostat.^65^ The production run was extended to 20 µs with a 20 fs integration time step.

### Atomistic molecular dynamics simulations

Atomistic molecular dynamics simulations of the MUT-16 FFR condensate were initiated from the final configuration of the corresponding Martini3-IDP^44^ coarse-grained trajectory. The coarse-grained system was converted to an all-atom representation using the initram.sh backmapping workflow, which automates the execution of backward.py to generate atom-istic coordinates and subsequently performs a sequence of energy minimization and position-restrained equilibration steps to relax the system.^66,67^ To reduce computational cost while preserving the condensate geometry and interfacial features, the simulation slab was trun-cated along the *z*-dimension by selectively removing solvent molecules, reducing the box size to 12 × 12 × 35 nm^3^. The resulting atomistic system comprised approximately 544,800 atoms. All atomistic simulations were performed using GROMACS (version 2018)^59^ with the Amber99sb-star-ildn-q^68–71^ force field in combination with the TIP4P-D^72^ water model. Following backmapping, the system was subjected to energy minimization using the steepest-descent algorithm to eliminate unfavorable contacts. The energy-minimized structure was equilibrated using a multistep protocol analogous to that employed for the coarse-grained simulations. First, the solvent and ions were equilibrated in the NVT ensemble while restrain-ing the protein coordinates. These restraints were then released, and additional equilibration was performed in the NVT ensemble to allow relaxation of the protein. Finally, the system was equilibrated under NPT conditions using a 2 fs time step. Temperature was maintained at 300 K using the Bussi-Donadio-Parrinello velocity-rescaling thermostat,^64^ and pressure was controlled at 1 bar using the Parrinello-Rahman barostat.^65^ Production simulations were carried out in the NPT ensemble for 1 *µ*s each across 10 independent replicas, yielding an aggregate sampling time of 10 *µ*s.

### Contact analysis

For contact analysis, we employed the "cascade computing" framework (https://github.com/comp-mol-biol/cascade_computing) previously developed in our lab. Designed for high-performance computing (HPC) clusters, this approach facilitates the calculation of pairwise contact data through a high degree of parallelization. It ini-tially stores pairwise residue contacts in an intermediate data structure, a so-called "contact record". Thus, the record provides a consistent yet resource-efficient starting point for spe-cific downstream analysis, eliminating the need for computationally expensive reprocessing of raw trajectories.

The parallelization strategy operates on two levels: first, the large molecular system is decomposed into numerous subsets of chain pairs assigned to independent computing nodes. Second, within each node, parallel Dask workers process individual chain pairs and trajectory segments to fully leverage CPU resources. Here we have used this approach to analyze a system of 2080 chain pairs with 48841 residue pairs each. Using one Dask worker per CPU (10 GB RAM/worker) across 40 nodes (40 CPUs each), we processed the full dataset (10 trajectories, 800 ns each, 100 ps resolution) in 15 hours without explicit fine-tuning. In a single run, we can derive contact matrices resolved by e.g. residue ID or amino acid type, for both contact frequency and lifetimes, solely from the contact record.

A residue pair is defined as being in contact if at least one corresponding atom pair is in contact. To efficiently exclude irrelevant chain pairs, we followed the approach of von Bülow et al.^73^ by first evaluating the distances between protein centers of mass (COM) before computing atomistic contacts. Furthermore, the analysis employs a dual-cutoff scheme consisting of a strict cutoff *d*, and a weaker cutoff *d*_lb_ = (*d* + Δ*d_ɛ_*). In a system with *N*_res_ residues, for any residue pair at position *x_i_, x_j_*at time *t_k_*to be in contact, the following condition must hold:

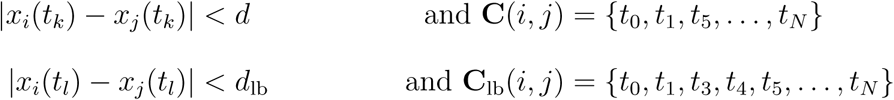

Based on the two contact sets **C** ⊂ **C**_lb_, which contain all timesteps where a pair is in contact, we process breaks in **C** in order to account for transient excursions. As Galvanetto et al. proposed, we additionally include respective frames from **C**_lb_ where the contact is maintained within the weaker contact cutoff *d*_lb_ before re-forming a strict contact.^20^ In this way, the lifetime of a contact initially formed below a cutoff of *d* is considered unbroken until the distance exceeds *d*_lb_. Specific parameters for the different contact analysis cases are provided in the Supplementary Information in Table S1.

The contact frequency *f* (*l, m*) for a selection of residues is calculated in (1) based on its number of frames in contact. We divide it by the number total contacts (*N_C_*) that form over the full length of the trajectory. More formally, for any aggregation of residue pairs *i, j* into sets *L, M* (e.g. amino-acids) we can map:

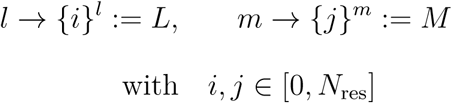

with *i, j* ∈ [0*, N*_res_] For the contact frequency it then holds:

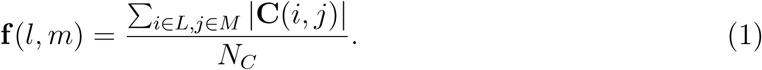

For the normalized contact frequency *f*^^^ we divide *f* by the abundance of each amino-acid pair. In this work, we extended the "cascade computing" framework to analyze specific non-covalent interactions, covering: (1) *π* − *π* stacking, (2) cation-*π* interactions, (3) salt bridges. We document specific bond types as an additional contact attribute in the contact record, based on the geometric definitions from Venkatakrishnan et al.^41^ Our implementation adopts the logic of their Python package getContacts, removing the dependence on VMD^74^ as a pre-requisite (https://github.com/getcontacts/getcontacts).

For the analysis of the hydrogen bonds we used MDAnalysis,^49,75^ in particular the corresponding HydrogenBondAnalysis class. (https://docs.mdanalysis.org/2.0.0/documentation_pages/analysis/hydrogenbonds. html).

### Ion and water analysis

We quantified the association of ions (Na^+^ and Cl^−^) and water with both amino-acid side chains and backbones within the MUT-16 FFR condensate. Ion–residue and water–residue contacts were defined using distance cutoffs of 4 Å for Na^+^, 5 Å for Cl^−^, and 4 Å for water molecules. These cutoffs were chosen to span the region between the first and second solvation shells, thereby capturing chemically meaningful coordination and hydrogen-bonding interactions without overcounting nonspecific solvent proximity.^76^.^77^ The protein–water interface was identified from one-dimensional density profiles computed along the *z*-axis, with the interface defined as the region where the protein density begins to decrease toward bulk solvent values. Density profiles of protein, water, and residue classes (acidic, basic, and polar) were calculated along the *z*-axis and averaged over the analyzed portions of all trajectories. Bridging interactions between similarly charged residues were identified by monitoring the simultaneous proximity of a counter-ion or water molecule to two residues of the same charge. For example, a Glu–Glu bridging event was defined when a Na^+^ ion or a water molecule was located within the respective distance cutoff of both residues, thereby mediating an otherwise electrostatically unfavorable interaction.

## Results

### Sequence composition reveals MUT-16 FFR as more representative of Human IDRs than FUS LCD

Because sequence composition and sequence patterning are key determinants of protein phase separation, we examined the MUT-16 FFR sequence and residue composition within the broader landscape of 28,058 human IDRs compiled by Tesei *et al.*,^10^ and in comparison to the canonical FUS LCD,^19,55^ using EBA^56^ scoring and RMSE of amino-acid composition vectors.

To quantify sequence similarity, a similarity score (EBA_min_) was computed for each human IDR relative to both MUT-16 FFR and FUS LCD. The resulting EBA_min_ distributions are summarized as box-and-whisker plots (Fig. 2A). Although the mean and median similarity scores are comparable for MUT-16 FFR and FUS LCD, their distributions differ markedly in spread. The interquartile range (25th–75th percentiles) of EBA_min_ values is broader for MUT-16 FFR, indicating greater variability in similarity to human IDRs. Consistently, the MUT-16 FFR distribution exhibits both higher upper-tail values (∼1.6 vs. ∼1.4) and lower minima than FUS LCD. Together, these features suggest that, despite similar average similarity, MUT-16 FFR spans a wider similarity landscape, reflecting greater sequence heterogeneity and reduced repetitiveness compared to the low-complexity profile of FUS LCD.

We next quantified compositional similarity by representing each human IDR as an amino-acid abundance vector and computing the root-mean-square error (RMSE) relative to the corresponding vectors of the MUT-16 FFR and the FUS LCD (Fig. 2B). The resulting RMSE distributions differ substantially between the two reference sequences. For the MUT-16 FFR, both the mean (∼ 12) and median (∼ 10) RMSE values are substantially lower than those obtained for the FUS LCD (mean ∼ 15, median ∼ 14). These trends indicate that, with respect to sequence composition, the MUT-16 FFR is more similar to a larger fraction of human IDRs than the FUS LCD.

To further elucidate the compositional similarity, we examined residue-abundance distributions across human IDRs and compared with those of MUT-16 FFR and FUS LCD (Fig. 2C). MUT-16 FFR contains all amino acids except Trp, which is itself rare in human IDR sequence analyzed,^10^ whereas FUS LCD is depleted in ten residue types (Cys, Glu, Phe, His, Ile, Lys, Leu, Arg, Val, and Trp). Notably, eleven residues in MUT-16 FFR fall within the interquartile range of residue abundance in of human IDRs, compared with only four in FUS LCD. Although both proteins are enriched in Gly, Gln, Ser, and Tyr, residues common in human IDRs, such as Lys and Leu, are reduced in MUT-16 FFR and absent in FUS LCD. These compositional differences account for the lower RMSE values of MUT-16 FFR and highlight its closer alignment with the global sequence landscape of human IDRs (Fig. 2B).

In conclusion, these analyses demonstrate that the MUT-16 FFR more closely resembles the broader class of human IDRs than the FUS LCD, supporting its relevance as a representative model system for studying the molecular principles underlying phase-separated protein condensates.

### MUT-16 FFR exhibits Upper Critical Solution Temperature (UCST) phase behavior

Given the high abundance of proline (Pro)^78^ and aromatic residues, we wondered whether MUT-16 FFR condensates are more stable at lower (20^◦^C) or elevated (40 ^◦^C) temperatures, and how this behavior compares to the reported disappearance of *Mutator foci* at higher temperatures *in vivo*,^51^ which we tested *in vitro*. Proline and aromatic-rich proteins such as the C-terminal domain of RNA Polymerase II, form more stable condensates at elevated temperatures and dissolve upon cooling,^79^ characteristic of Lower Critical Solution Temperature (LCST) phase behavior. Other proteins such as FUS LCD exhibit Upper Critical Solution Temperature (UCST) behavior.^25,80^ We observed the phase separation of MUT-16 FFR at 20^◦^C, 30^◦^C and 40^◦^C in experiments, where we controlled the temperature as described re-cently by Fritsch et al.^58^ Encapsulating the protein solution makes it possible to quantify the volume of the dense phase in comparison to the total volume of the solution confined by the oil droplets. The ratio of these volumes defines the volume fraction (Fig. 3A). As expected, higher protein concentration resulted in an increase of the volume of droplets (Fig. 3B), with experiments conducted at 10 *µ*M, 15 *µ*M and 20 *µ*M. When the volume fraction was normalized to its value at 20^◦^C for each protein concentration, it showed that increasing the temperature reduces the tendency to phase separate, as evidenced by a reduced volume fraction (Fig. 3C). At an intermediate temperature (30^◦^C), the normalized volume frac-tion decreases by approximately a factor of two as the concentration increases from 10 *µ*M to 15 *µ*M, and further decreases by a similar factor upon increasing the concentration to 20 *µ*M. At a higher temperature (40^◦^C), the volume fraction becomes negligible at 15 *µ*M and remains negligible at 20 *µ*M. These observations demonstrate a strong and non-linear dependence of condensate volume fraction on both temperature and protein concentration. Overall, the *in vitro* measurements provide compelling evidence that the MUT-16 FFR ex-hibits UCST phase behavior, as demonstrated by the pronounced temperature-dependent modulation of condensate formation, where phase separation is favored at lower temperatures and progressively suppressed upon heating.

Previous studies define UCST sequence heuristics based on five parameters: length, charge content (R+K+D+E), zwitterionic character [(R+K)/(R+K+D+E)], aromatic con-tent (Y+H+W+F), and Arg enrichment [R/(R+K)], with thresholds of *>* 100 residues, *>* 20% charge, 45–55% positive charge fraction, *>* 10% aromatic content, and *>* 85% Arg enrichment.^81^ MUT-16 FFR satisfies the length (172 residues), aromatic content (20.35%), and Arg enrichment (85.71%) criteria. Its charge content (10.47%) and charge balance (38.89% positive charge fraction) are slightly below the nominal thresholds; however, previ-ous studies show that resilins exhibiting UCST behavior typically contain 12–15% charged residues,^82^ indicating that such values remain compatible with UCST. Similarly, the deviation in charge balance corresponds to ∼ ±7% from ideal zwitterionic character, which is close to the expected ±5% window.^81^ Overall, these features indicate that MUT-16 FFR is consistent with UCST-like sequence characteristics and rationalize our *in vitro*, where MUT-16 FFR phase separation was reduced as temperature increased, and previous *in vivo* observations,^51^ where *Mutator focii* disappeared as temperature was raised.

Considering the sequence comparison, the similarities of temperature response, but also the previously reported similar effect of truncations of the MUT-16 sequence *in vitro* and *in vivo*,^39,51^ we conclude that studying MUT-16 condensates on their own can elucidate drivers of *Mutator focii* condensation *in vivo*.

### Molecular dynamics simulations highlight frequent contacts of polar and aromatic groups as well as charge-charge interactions

Having established MUT-16 FFR as a model system for phase-separated condensates, we turned to atomistic molecular dynamics simulations to elucidate how interactions among different amino acids shape MUT-16 condensates (Movie S1). Ten atomistic trajectories (1 *µ*s) were initiated by reconstructing atomistic detail from an initial coarse-grained simulation where MUT-16 FFR spontaneously phase-separated (Movie S2). We characterized the interaction dynamics of the phase-separated MUT-16 FFR condensate at atomistic resolution by analyzing both unnormalized and abundance-normalized contact frequencies, followed by quantification of contact lifetimes (Fig. 4). All interaction analyses were performed at the level of amino acid side chains, to capture the chemical diversity and interaction specificity within the condensate. Together, these metrics enable a complementary view of interaction frequency, compositional bias, and dynamics within the condensate.

**Figure 4:**
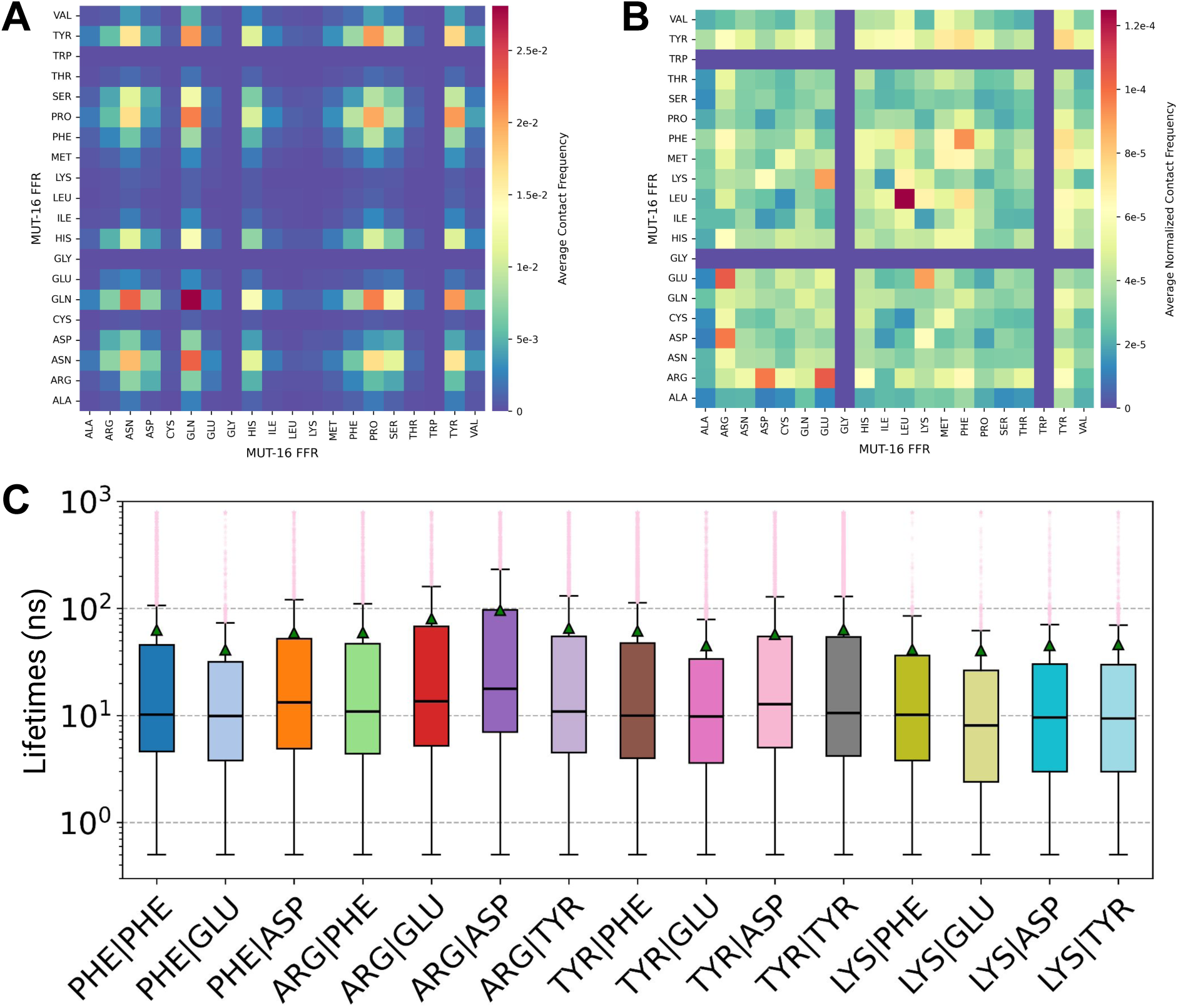
Residue-resolved contact frequencies and persistence of side-chain interactions in the MUT-16 FFR condensate. **A, B.** Residue resolved contact maps derived from atomistic simulations of the MUT-16 FFR condensate, shown in both unnormalized and abundance-normalized heatmaps. The maps report contact frequencies for side chain-side chain (SC:SC) interactions between all amino acid pairs, where unnormalized frequencies reflect the raw occurrence of contacts, and abundance-normalized frequencies account for sequence composition effects to highlight intrinsically preferred interactions. **C.** Box-and-whisker plots of lifetimes for SC:SC contacts between amino-acid pairs. The central line indicates the median, the green triangle denotes the mean, and high-persistence outliers beyond the upper tail are shown as translucent pink stars.

The unnormalized interaction profile reveals a high frequency of contacts between polar uncharged residues, particularly Gln and Asn (Fig. 4A). This enrichment reflects their high abundance in the MUT-16 FFR sequence (Fig. 2C) and the ability of their side-chain amide groups to both donate and accept hydrogen bonds. In addition, Tyr and Pro, which are also abundant in the MUT-16 FFR sequence, exhibit frequent interactions with other residues of the same type, strong mutual interactions, and extensive contacts with other residues, consistent with previous studies^27,79^ that highlight their central roles in mediating interac-tions. Furthermore, despite being as abundant as Tyr (Fig. 2C), Ser shows significantly fewer interactions, highlighting the importance of the Tyr phenyl ring in favoring interaction.

By contrast, the abundance-normalized contact map reveals a distinct interaction pattern, which emphasizes specific residue–residue interaction propensities rather than overall residue abundance.^38^ Hydrophobic Leu–Leu interactions emerge as the most prominent contacts within the MUT-16 FFR condensate (Fig. 4B), reflecting putatively favorable interactions of their aliphatic side chains. Note that relatively few such events are observed, which is not surprising considering their low abundance, with a single Leu residue in the MUT-16 FFR sequence (Fig. 2C). These are followed by Arg–Glu and Arg–Asp interactions, consistent with favorable salt-bridge formation between oppositely charged guanidinium and carboxylate groups, with Arg–Glu contacts occurring more frequently than Arg–Asp. Phe–Phe interactions are moderately strong and more prominent than Tyr–Tyr contacts, which may indicate that local sequence context favors interchain interactions of the five Phe residues. Despite this, Tyr residues participate in a wide range of interactions with multiple residue types, indicative of a promiscuous interaction profile. Notably, Arg–Glu and Arg–Asp interactions are substantially more prevalent than Lys–Glu contacts, which, although capable of forming salt bridges, display only moderate interaction propensity; Lys–Glu interactions nevertheless exceed those of Lys–Asp. Trp does not contribute to the contact network because it is absent from the MUT-16 FFR sequence, whereas Gly exhibits negligible interactions owing to its minimal side chain. Other aliphatic side chains of Ala and Val exhibit uniformly weak interactions across all partners.

In conclusion, unnormalized and abundance-normalized contact maps show that interactions within the MUT-16 FFR condensate are governed by both polar and charged residues. Polar residues (Gln and Asn) contribute through high sequence abundance, whereas charged residues (Arg, Glu, and Asp) contribute through their strong intrinsic interaction strength. Tyr residues are abundant in the sequence and exhibit correspondingly high contact frequencies, whereas Phe residues, despite their lower abundance, display disproportionately high contact frequencies relative to their abundance. Together, these results elucidate the interplay between prevalence and propensity in governing condensate interaction dynamics.

### Contact lifetimes correlate with normalized, but not unnormalized, contact frequency

To probe contact persistence, we quantified side-chain contact lifetimes and correlated mean lifetimes with unnormalized and abundance-normalized contact frequencies. Lifetime distributions for residue pairs with high normalised contact frequencies (Fig. 4B) are shown as box-and-whisker plots in Fig. 4C, while mean lifetimes for all residue pairs are summarized in Fig. S2B. The temporal resolution of the analysis is limited by the 0.1 ns sampling interval, resulting in a lower tail fixed at 0.1 ns. We consider fully persistent contacts across the entire trajectory, allowing for potential interplay between short- and long-lived contacts. To account for outliers that substantially affect the mean lifetimes, we present the full lifetime distribution and discuss the corresponding medians.

Overall the lifetimes of contacts are short, but there are a few long-lived contacts. Depending on the residue pair in the MUT-16 FFR atomistic simulations, between 10^2^ and more than 10^5^ binding events were observed (Fig. S2A). Across selected interaction types, mean lifetimes substantially exceed the corresponding medians, indicating strongly right-skewed, non-Gaussian lifetime distributions dominated by rare long-lived events (Fig. 4C). In many cases, the influence of these events is sufficient for mean values to approach or slightly exceed the upper-quartile values, with upper tails extending to approximately ∼ 100 ns across residue pairs. Consistent with this behavior, the lifetime histograms exhibit a sharp decay beyond ∼ 100 ns, indicating that long-lived contacts are generally infrequent (Fig. 5A). Interestingly, for specific residue pairs such as Arg-Asp and Arg-Glu, approximately one in ten contacts persists longer than 100 ns (Fig. 5A). Median lifetimes for most interactions cluster around ∼ 10 ns, indicating a broadly similar timescale of contact persistence across residue pairs for the pairs shown in (Fig. 5C). Considering the contacts of all 19 amino acid types in the FFR sequence and computing the lifetimes of their pairwise contacts, we obtain a median of ∼ 9.8 ns. Notably, interactions involving Arg and Asp residues exhibit higher median lifetimes, generally exceeding 10 ns on average (Fig. S3), reflecting their enhanced stability. In contrast, Lys-mediated interactions represent a clear exception, displaying comparatively lower median lifetimes of approximately ∼ 8 ns (Fig. 4C, S3), suggesting reduced contact persistence relative to other charged residues.

**Figure 5:**
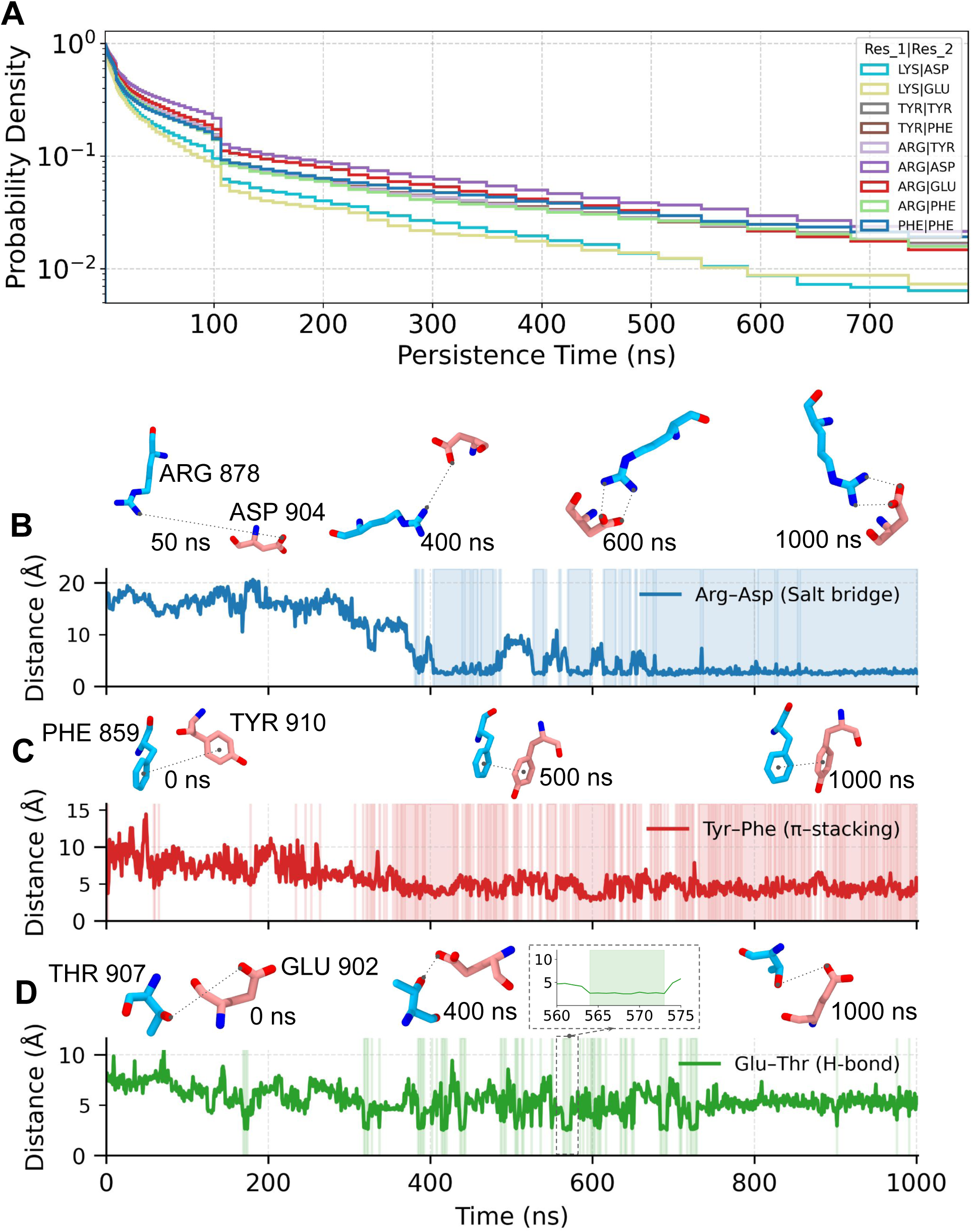
Representative persistence time distributions and interaction dynamics of side-chain contacts. **A.** Probability density distributions of lifetimes for selected amino-acid pairs. **B.** Distance time series for a representative Arg–Asp salt bridge; shaded regions indicate bound states defined by a 4 Å donor–acceptor cutoff. **C.** Distance time series for a representative Tyr–Phe *π*–*π* interaction; bound states are defined by a 5 Å center-of-mass cutoff between aromatic rings. **D.** Distance time series for a representative Thr–Asp hydrogen bond; bound states are defined by a 3.5 Å donor–acceptor cutoff. Representative snapshots corresponding to selected time points are shown above each time series.

The upper-quartile and upper-tail lifetimes are distinct across residue pairs (Fig. 4C). Arg–Asp interactions exhibit the largest upper-quartile values (∼ 80 ns), followed by Arg–Glu (∼ 60 ns). Interestingly, although abundance-normalized contact frequencies indicate stronger Arg-Glu interactions than Arg-Asp (Fig. 4B), the longer mean persistence of Arg-Asp contacts (Fig. S2B) suggests that interaction prevalence and lifetime are not strictly correlated (Fig. 4C). We ensured that this disparity is not driven by a few rare, long-lived contacts by analyzing individual Arg-Asp and Arg-Glu pairs in the MUT-16 FFR sequence (Fig. S4), and observed that Arg-Asp pairs generally exhibit higher median and mean lifetimes. This disparity may reflect differences in side-chain geometry: the longer, more flexible side chain of Glu likely facilitates more frequent encounters with Arg but also leads to greater fluctuations in the contact profile. In contrast, the shorter side chain of Asp requires interaction partners to approach more closely, resulting in fewer contacts but with longer lifetimes.

Further, Lys–Asp and Lys–Glu interactions display substantially lower upper-quartile values (∼ 35 ns). Aromatic interactions exhibit distinct statistical signatures: Tyr–Tyr interactions show a broader interquartile range than Phe–Phe, while Arg–Tyr and Arg–Phe interactions display greater variability compared to Lys-based contacts. Aromatic interactions (Tyr–Tyr, Tyr–Phe, and Phe–Phe) also exhibit the highest density of high-persistence outliers, followed by Arg-mediated interactions, whereas Lys-based interactions show comparatively few outliers. The scarcity of long-lived Lys-mediated contacts is further evident from their low probability density at long times (Fig. 5A).

Mean lifetimes (Fig. S2B) show no apparent correlation with unnormalised contact frequency (Fig. S2C), whereas a moderately strong correlation is observed with abundance-normalised contact frequency (Fig. S2D). Within this framework, Leu-Leu interactions emerge as among the most persistent contacts in the MUT-16 FFR condensate. Although this trend is consistent with their high abundance-normalized contact frequency (Fig. S2D), the relatively limited number of binding events (10^2^–10^3^; Fig. S2A) suggests that the el-evated lifetimes may be partially influenced by statistical sampling. Salt-bridge-like Arg-Asp and Arg-Glu interactions exhibit high mean lifetimes that correlate strongly with their abundance-normalized contact frequencies (Fig. S2D), consistent with the intrinsic stability of guanidinium-carboxylate salt bridges. Furthermore, Pro-Pro and Pro-Tyr contacts exhibit mean lifetimes of approximately 70 ns and 60 ns, respectively (Fig. S2B). Despite the enrichment of Pro residues in MUT-16 FFR and the large number of Pro-mediated contacts formed in the system (Fig. 4A, Fig. S2C), abundance normalisation reveals that Pro-based interactions occur less frequently than many other contact types (Fig. 4A, Fig. S2D). Many Pro-based interactions (Pro-Pro, Pro-Tyr,^79^ Pro-Leu, and Pro-Phe) display relatively long lifetimes (Fig. S2B, Fig. S2C), suggesting that these contacts may nonetheless contribute to stabilising the MUT-16 condensate, consistent with previous studies highlighting the role of Pro-mediated interactions in phase-separated systems.^79^ Finally, the correlation analysis reveals that highly abundant residues in MUT-16 FFR, such as Gln and Asn, engage in a large number of contacts (Fig. 4A; Fig. S2C), yet the vast majority of these interactions are short-lived underscoring the distinction between contact abundance and interaction stability.

To illustrate representative interaction dynamics (Fig. 5), we analyzed distance time series for residue pairs capable of forming salt bridges (Arg878–Asp904), *π*–*π* stacking (Phe869-Tyr910), and hydrogen bonds (Thr907–Glu902). Distance-based cutoffs were applied to define bound residue pairs: 4 Å for pairs capable of forming salt bridges, 6 Å for pairs capable of engaging in *π*-*π* stacking interactions, and 3.5 Å for pairs capable of forming hydrogen bonds. For the Arg–Asp pair, the inter-residue distance initially remains large (∼ 20 Å), decreases gradually after ∼ 200 ns, and forms the first contact between ∼ 400 and 500 ns, followed by repeated dissociation and association events (Fig. 5B). After ∼ 600 ns, the interaction becomes more stable, exhibiting reduced fluctuations, as stable interactions is formed as illustrated by the conformations in (Fig. 5B). By contrast, the Tyr–Phe pair begins at a shorter separation (∼ 10 Å) and forms a stable contact around ∼ 350 ns (Fig. 5C), which persists for extended periods but with frequent transient disruptions. Finally, the Thr–Glu interaction is highly dynamic and short-lived. The inter-residue distance initially remains at ∼ 8 Å and exhibits transient hydrogen-bond formation between approximately 400 and 700 ns before dissociating completely (Fig. 5D). Individual contact events persist for only ∼ 5–10 ns, as illustrated in the zoomed-in region between 560 and 575 ns. Together, these examples indicate how a weak hydrogen bonding contact can be broken very quickly and that some salt-bridge interactions and aromatic-mediated contacts can persist for long-times in some cases, while the typical such contact is still very short lived (Fig. 4C and Fig. S3).

We conclude that most interactions are highly dynamic, particularly those involving the abundant polar uncharged residues Asn and Gln. The contacts of aromatic side chains and charged residues are also almost always short-lived, but a few rare contacts persist over extended durations. The contact lifetimes do not correlate with the absolute frequency of contacts, as these are dominated by weakly interacting but highly abundant residues, but with the contact frequency normalized by residue-pair abundance.

### Fast dynamics of hydrogen-bonds, salt-bridges, cation–*π*, and *π*–*π* stacking

To further characterize the interactions of different residues in MUT-16 FFR condensates we followed specific noncovalent interactions within the condensate, including hydrogen bonds, salt bridges, cation–*π*, and *π*–*π* stacking, by comparing unnormalized and abundance-normalized contact frequency/fractions together with their corresponding lifetimes.

Analysis of hydrogen bonding interactions reveals that side chain-side chain contacts contribute the largest number of hydrogen bonds within the MUT-16 FFR condensate, followed by backbone-backbone interactions, with backbone-side chain interactions contributing the fewest (Fig. S5A). We further investigated the side chain-side chain hydrogen bonds. Residue-abundance normalized contact maps reveal Arg–Glu interactions as the most prevalent, followed closely by Arg–Asp, consistent with favorable hydrogen bonding between guanidinium and carboxylate functional groups (Figure 6A). By contrast, the unnormalized contact maps show a significantly higher frequency of Arg–Asp interactions relative to Arg–Glu (Figure S5B), reflecting the underlying sequence composition. Normalization also highlights a significant contribution from Lys interactions with negatively charged residues (Figure 6A), which are largely obscured in the unnormalized analysis (Figure S5B). In addition, the un-normalized contact maps indicate a high prevalence of hydrogen bonds between Asn and Gln (Figure S5B), consistent with their high abundance in the sequence; a representative Asn–Gln hydrogen bond is shown in Figure S5C. Polar residues such as Ser also contribute appreciably to the hydrogen-bond network in both unnormalized and abundance-normalized contact maps.

**Figure 6:**
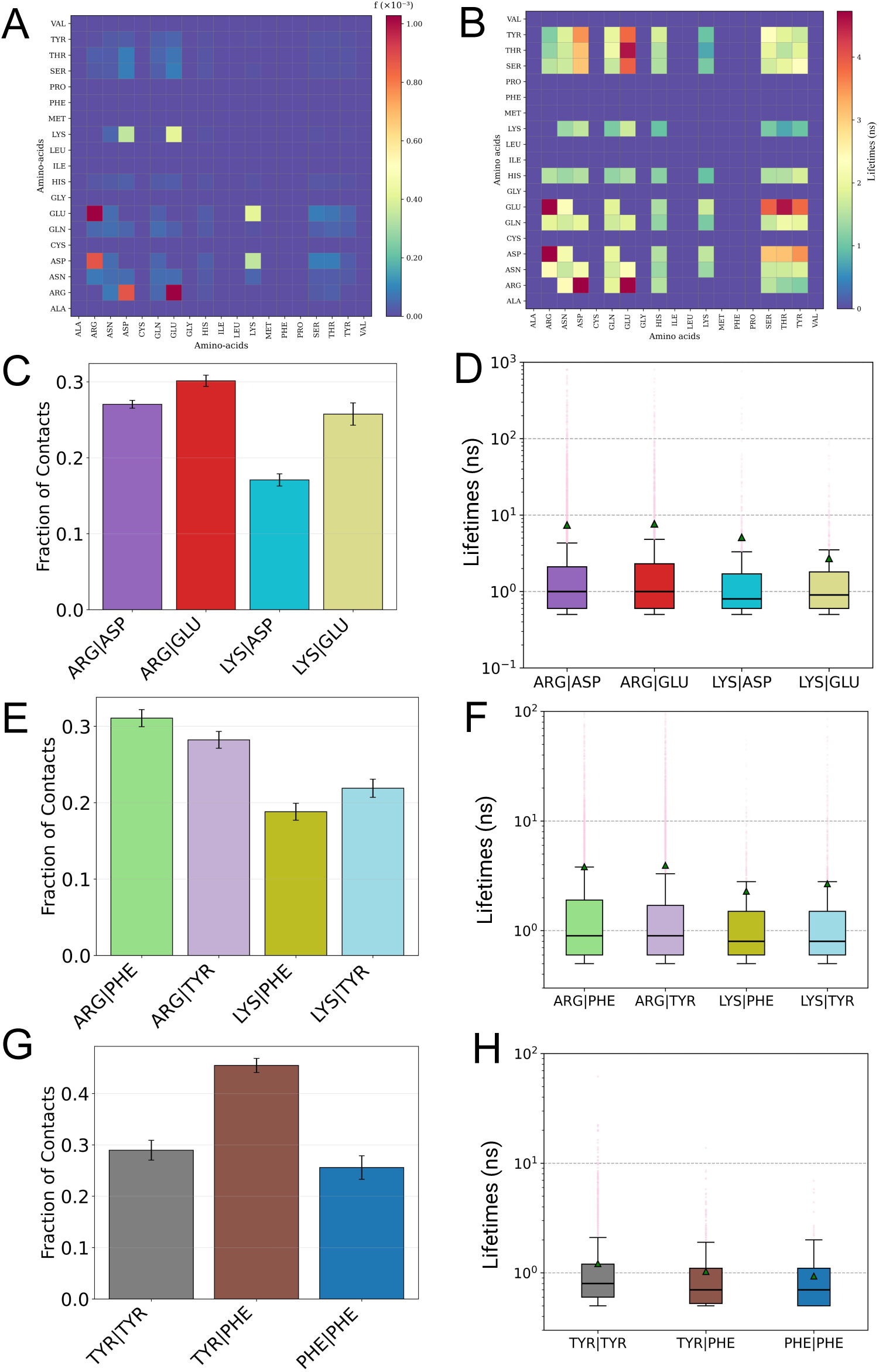
Distinct noncovalent interaction modes and their persistence in the MUT-16 FFR condensate. **A,B.** Hydrogen-bond interactions in the MUT-16 FFR condensate: (A) heatmap of contact frequencies and (B) mean lifetimes. **C,D.** Salt-bridge interactions: (C) fraction of residue-residue contacts forming salt bridges, indicating the proportion of all salt bridges in the system contributed by each residue pair; and (D) box-and-whisker distributions of salt-bridge lifetimes. **E,F.** Cation–*π* interactions: (E) fraction of contacts and (F) persistence-time distributions. **G,H.** *π*–*π* stacking interactions: (G) fraction of residue-residue contacts, indicating the contribution of each residue pair and (H) persistence-time distributions.

Lifetime analysis of hydrogen bond contacts closely mirrors the trends observed in the abundance-normalized contact maps. Arg–Glu and Arg–Asp interactions exhibit the longest lifetimes, with mean lifetimes of approximately ∼ 5 ns (Figure 6B). Notably, hydrogen bonds formed between acidic residues (Glu or Asp) and hydroxyl-containing residues such as Thr, Ser, and Tyr are more persistent than those involving Lys–Glu or Lys–Asp pairs, despite the latter being capable of forming salt bridges. This behavior likely reflects the greater directionality and geometric specificity of hydroxyl-mediated hydrogen bonding compared to the more flexible ammonium–carboxylate interactions of Lys. Interactions involving Gln and Asn also contribute to the hydrogen-bond network, consistent with their pronounced enrichment in the MUT-16 FFR sequence (Figure 2C); however, these contacts are generally weaker and more transient than the salt-bridge-associated hydrogen bonds (Figure 6B).

We further analyzed salt-bridge interactions by quantifying contact fractions using both unnormalized and abundance-normalized metrics and comparing these trends with the corresponding lifetime distributions. In the abundance-normalized analysis, the ranking of salt-bridge contact fractions (Arg–Glu > Arg–Asp > Lys–Glu > Lys–Asp; Figure 6C) closely mirrors that observed for normalized hydrogen-bond contacts. By contrast, the unnormalized analysis reveals a significantly different pattern, with Arg–Asp contacts occurring at more than twice the frequency of Arg–Glu contacts, which in turn exceed Lys-based interactions by more than a factor of two (Figure S5D), reflecting contributions beyond sequence abundance alone, given that Asp is not twice as abundant as Glu. Persistence-time analysis reveals comparable median lifetimes across all salt bridges (∼ 1 ns); however, Arg-mediated interactions exhibit broader interquartile ranges and higher mean lifetimes (∼ 10 ns) than Lys-based contacts (∼ 5 ns). Notably, Arg forms some long-lived, salt bridges with lifetimes beyond 10 ns or even 100 ns, with some salt-bridges persisting for almost the entire duration of the trajectory (Figure 6D). Such long-lived events, with peristence times beyond 100 ns are rarely observed for Lys. Lys-Asp and Lys-Glu interactions display similar medians, interquartile ranges, and mean lifetimes.

Analysis of cation-*π* interactions shows that Arg-mediated contacts are consistently more prevalent than Lys-mediated contacts in both unnormalized (Figure S5E) and abundance-normalized (Figure 6E) contact fractions. This difference is consistent with the ability of the guanidinium group of Arg to engage in multivalent interactions with aromatic residues, combining cation-*π* interactions, hydrogen bonding, and sp^2^-*π* interactions, capabilities that are absent for the aliphatic ammonium group of Lys.^35,39,83–86^ Lifetime analysis reveals that all cation-*π* interactions are short-lived, with subnanosecond median lifetimes. Mean lifetimes are ∼ 3 ns for Arg-Tyr and ∼ 2 ns for Arg-Phe (Figure 6F). Although Lys-Tyr interactions display interquartile ranges and mean values comparable to Arg-Tyr and Arg-Phe, they exhibit markedly fewer long-lived outliers, whereas Arg-mediated interactions populate the upper tail of the distribution with rare events extending to ∼ 100 ns. In contrast, Lys-Phe interactions show a substantially narrower interquartile range and lack outliers beyond ∼ 10 ns. Interestingly, cation-*π* interactions formed by both Arg and Lys are significantly less persistent than the corresponding salt-bridge interactions, underscoring their more transient contribution to condensate stabilization.

Finally, we examined *π*–*π* stacking interactions. In the unnormalised contact fractions, Tyr-Tyr interactions are most prevalent, followed by Tyr-Phe and Phe-Phe (Figure S5E), reflecting the higher abundance of Tyr in the MUT-16 FFR sequence. Upon abundance normalisation, this trend no longer holds, as Tyr–Phe interactions are significantly stronger than Tyr–Tyr and Phe–Phe interactions (Figure 6G). Lifetime analysis shows that *bona fide π*–*π* stacking interactions are highly transient, with median lifetimes below 1 ns and narrow interquartile ranges (Figure 6H). In contrast, when aromatic interactions are defined using distance-only criteria without angular constraints, the median persistence time increases to approximately 10 ns (Figure 4C). To examine the role of angular constraints more closely, we analyzed a representative Tyr-Phe pair that appears to maintain prolonged contact based on distance alone (Fig. S6). However, applying angular constraints reveals that true *π*–*π* stacking occurs only rarely, with the interaction frequently fluctuating between bound and unbound states. Together, these observations underscore the importance of geometric alignment in defining *π*–*π* stacking interactions^27^ and show that proximity alone may substantially overestimate their persistence.

Overall, different non-covalent interactions give rise to short-lived contacts in the MUT-16 condensates. Some longer-lived interactions are found, for instance, for Arg-mediated salt-bridge and cation-*π* interactions (Movie S1), as well as Tyr-mediated cation-*π* and *π*–*π* stacking interactions.

### Na^+^ ions associate with and bridge negatively charged residues to regulate MUT-16 FFR dynamics

Ions such as Na^+^ and Cl^−^ play a critical role in modulating biomolecular phase separation by screening electrostatic interactions and mediating residue–residue contacts. The dense phase of MUT-16 is somewhat depleted in Na^+^ and mostly devoid of Cl^−^ ions compared to the aqueous phase (Fig. S7). The concentration profile of Na^+^ tracks the concentration profile of the acidic residues, which suggest that negatively charged side chains are interacting with Na^+^. To better understand the interactions of ions with the MUT-16 FFR condensate, we quantified the average number of Na^+^ and Cl^−^ ions interacting with each amino acid (Fig. 7). We next analyzed the lifetimes of ion–side-chain interactions to assess the stability and lifetime of these associations (Fig. 8). Finally, we investigated the role of counter-ions in mediating bridging interactions between pairs of similarly charged residues (Fig. 8), such as Glu-Glu or Arg-Arg, that would otherwise experience electrostatic repulsion and interact only weakly in the absence of ions. Ion-residue interactions were quantified using distance cutoffs of 4 Å for Na^+^ and 5 Å for Cl^−^ (Fig. 7). These cutoff values were chosen with reference to the reported positions of the first and second hydration shells of the respective ions,^76^ selecting intermediate distances to avoid being overly restrictive or overly permissive in defining ionresidue contacts. Interaction counts were averaged over 10 independent replicas, and the standard error of the mean was estimated from the replica-level averages.

**Figure 7:**
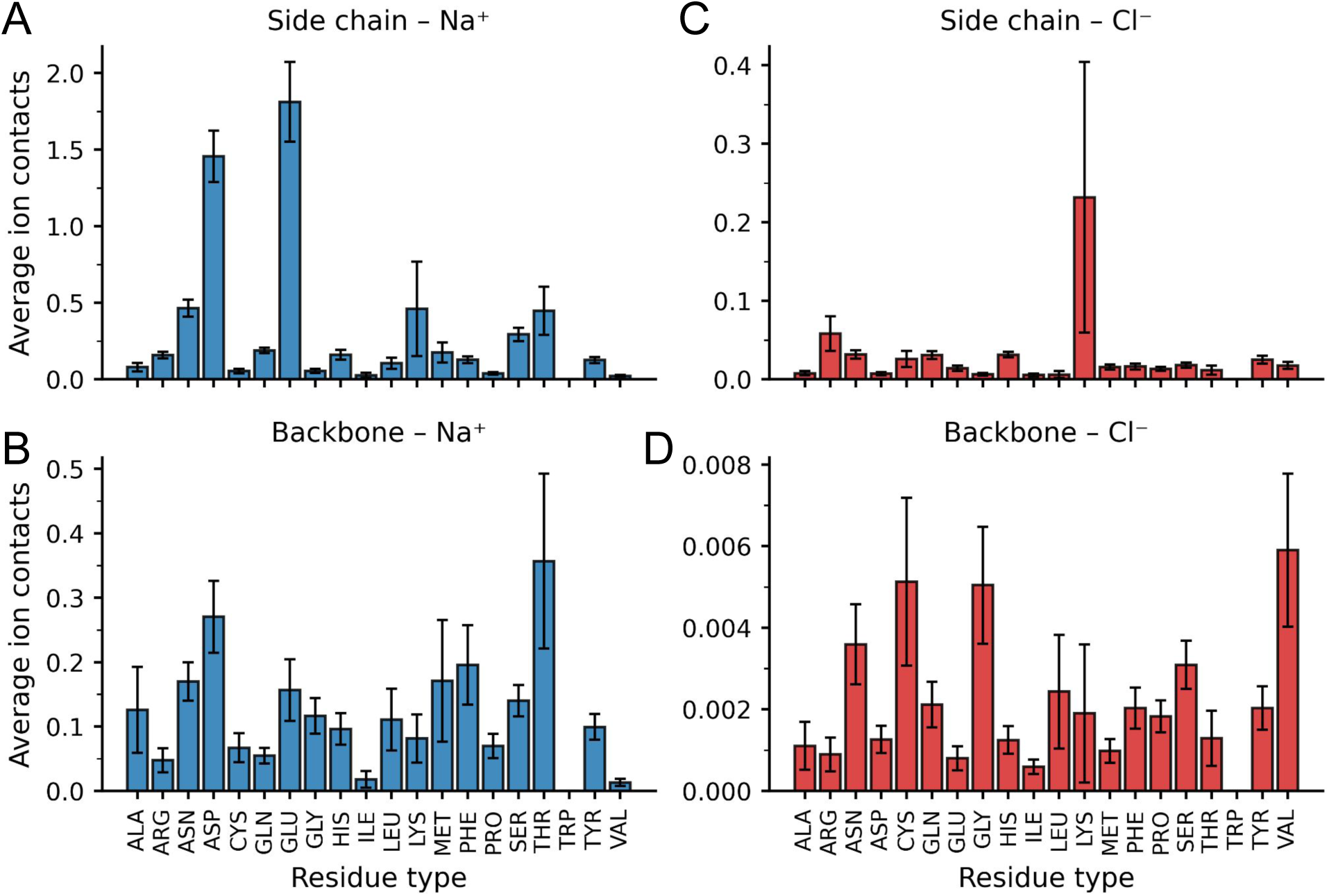
Residue-resolved ion interactions within the phase-separated MUT-16 FFR condensate. **(A,B)** Average number of Na^+^ ions interacting with amino-acid side chains (A) and backbones (B), highlighting preferential association of sodium with negatively charged and polar residues. **(C,D)** Average number of Cl^−^ ions interacting with amino-acid side chains (C) and backbones (D), showing substantially weaker interactions relative to Na^+^. Ion–residue interactions were defined using distance cutoffs of 4 Å for Na^+^ and 5 Å for Cl^−^. Values represent averages over 10 independent replicas, with error bars indicating the standard error of the mean.

**Figure 8:**
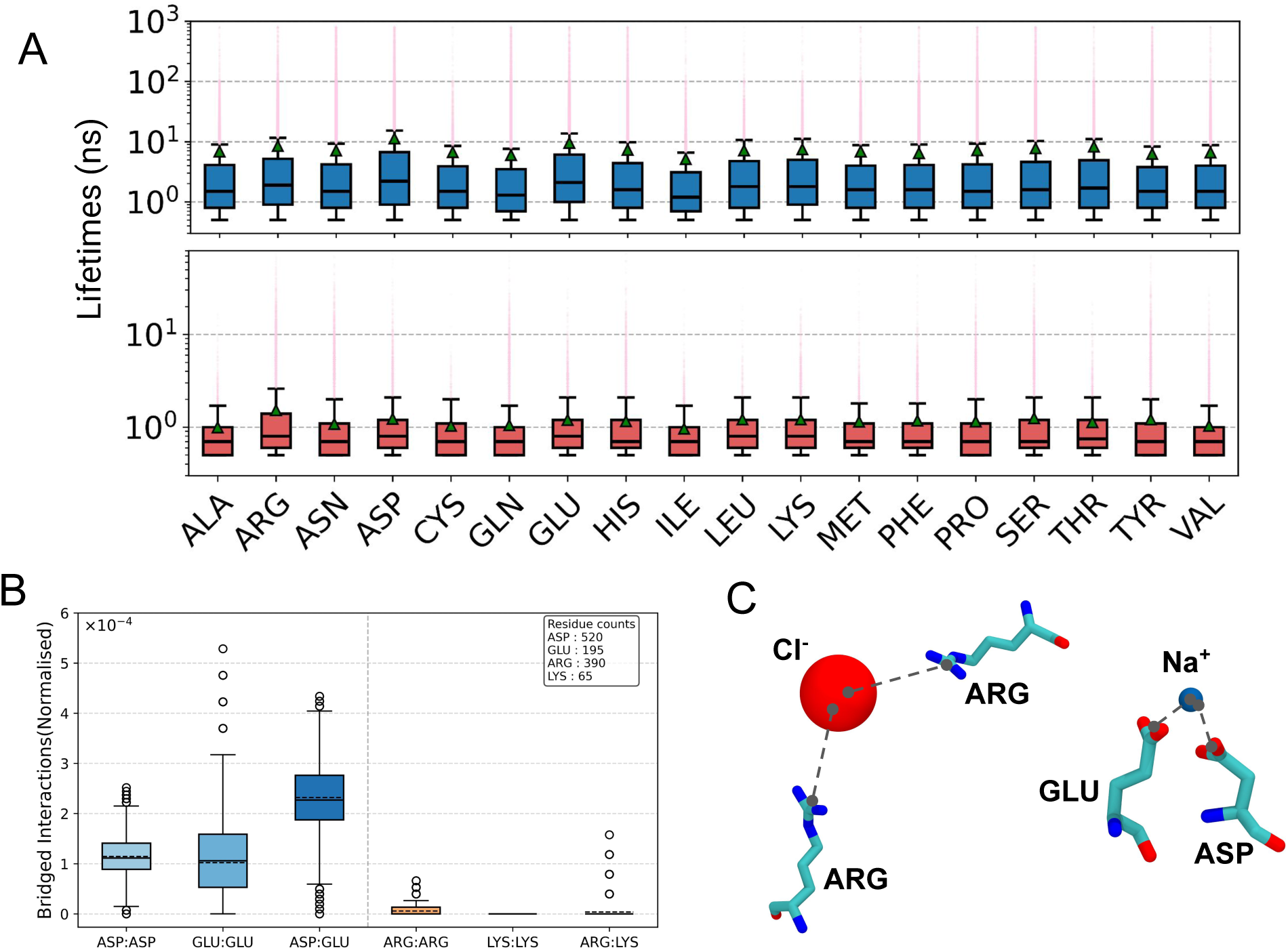
Ion-mediated interactions within the MUT-16 FFR condensate. **(A)** Box-and-whisker plots showing the lifetimes of ion–residue side-chain interactions. Distributions for Na^+^ interactions are shown in the upper panel (blue), whereas distributions for Cl^−^ interac-tions are shown in the lower panel (red). **(B)** Frequency of ion-bridged interactions between similarly charged residue pairs, normalized by amino-acid pair abundance. **(C)** Represen-tative snapshots illustrating Cl^−^- and Na^+^-mediated bridging interactions between Arg–Arg and Glu–Asp side chains, respectively.

Na^+^ ions exhibit strong and preferential interactions with negatively charged residues within the MUT-16 FFR condensate. Among these, Glu displays the highest affinity, interacting with approximately ∼ 2 Na^+^ ions per residue on average (Fig. 7A), followed by Asp, which coordinates roughly ∼ 1.5 Na^+^ ions per residue. Carboxylate side chains are known to bind multiple Na^+^ ions by bridging them through oxygen atoms, forming bidentate coordination motifs or “chelate rings”, as has been established in investigations of physical chemistry of interactions of acidic side chains and cations.^33^ Glu in the MUT-16 FFR condensate ex-hibits a higher propensity to engage in bidentate coordination with Na^+^ compared to Asp. While Glu dominates in side-chain-mediated interactions, backbone association with Na^+^ is more pronounced for Asp, with an average of ∼ 0.3 ions per residue compared to ∼ 0.2 for Glu (Fig. 7B). This trend is consistent with prior work on the interactions of Asp and Glu, showing that the shorter side chain of Asp confines carboxylate groups to a smaller effective volume, thereby increasing local charge density and drawing counter ions closer to the backbone region.^32^ Consistent with this mechanism, we observe instances in which a single Na^+^ ion bridges interactions between the Asp side chain and backbone; a representative example is shown in Fig. S9A.

Several neutral and polar residues exhibit moderate Na^+^ association, with Asn and Thr each interacting with approximately ∼ 0.5 Na^+^ ions per residue on average (Fig. 7A), likely mediated by coordination to side-chain carbonyl or hydroxyl oxygen atoms. Notably, Thr exhibits the highest level of backbone-associated Na^+^ interactions (Fig. 7B). This behavior likely arises from cooperative coordination involving backbone carbonyl oxygens in proximity to the polar side chain, as observed in the simulations (Fig. S9B). As a result, the total Na^+^ association of Thr (bb + sc) ranks third overall, following Glu and Asp, and exceeds that of Asn (Fig. S8A). Interestingly, Lys also shows appreciable Na^+^ association (∼ 0.5 ions per residue) despite its positive charge, likely arising from indirect interactions mediated by local electrostatic environments and nearby counter-ions, as observed in the simulations (Fig. S9C).

By contrast, interactions with Cl^−^ ions are markedly weaker overall. Lys is the most strongly interacting residue, averaging ∼ 0.22 Cl^−^ ions, albeit with a large statistical uncertainty , followed by Arg with fewer than 0.1 Cl^−^ ions on average (Fig. 7B). The relatively low level of Cl^−^ association is likely due to the overall negative charge of the condensate (net charge of −4 per chain, corresponding to an NCPR of −0.023 per sequence), which sums to −260 for 65 chains. This net negative charge disfavors the accumulation of negatively charged ions while promoting the enrichment of Na^+^ (Fig. S7). Backbone-resolved analysis indicates that Val, Cys, and Gly backbones recruit the largest number of Cl^−^ ions (Fig. 7B). These residues possess small or nonpolar side chains that leave the backbone more solvent-exposed, facilitating weak electrostatic and hydrogen-bond-like interactions between Cl^−^ ions and the partially positive backbone amide NH groups.^31^

We next analyzed the lifetimes of ion–residue side-chain interactions, as showing with a box-and-whisker plot in Figure 8A. Na^+^ interactions display broader lifetime distributions than Cl^−^, with median lifetimes of ∼ 1 ns compared to subnanosecond medians for Cl^−^. Consistent with their strong electrostatic affinity, Asp and Glu exhibit the highest mean Na^+^ lifetimes (∼ 10 ns), and Na^+^ interactions with Asp, Glu, Asn, and Gln feature rare, trajectory-spanning outliers. In contrast, Cl^−^ interactions are generally shorter lived. Among basic residues, Arg shows the longest-lived Cl^−^ associations, with outliers extending to ∼ 80 ns, whereas Lys–Cl^−^ interactions exhibit substantially shorter upper tails, with outliers limited to ∼ 30 ns. These trends indicate that ion-specific chemistry and side-chain functionality strongly modulate both the stability and lifetime heterogeneity of ion–residue contacts within the condensate.

We further investigated whether counter-ions mediate bridging interactions between pairs of similarly charged residues within the condensate (Figure. 8C). To this end, we quantified the total number of ion-bridged residue–residue interactions, normalized by the abundance of each residue pair (Fig. 8B). We find that Na^+^-mediated bridging between negatively charged residues is substantially more prevalent than Cl^−^-mediated bridging between positively charged residues, likely reflecting the limited presence of Cl^−^ ions within the negatively charged MUT-16 FFR condensate (Fig. S7). Among Na^+^-mediated bridges, Asp–Glu interactions occur more frequently than either Asp–Asp or Glu–Glu interactions. Although Glu–Glu and Asp–Asp bridges occur with similar median frequencies, Glu–Glu interactions exhibit a broader interquartile range (Q1–Q3) and a larger upper quartile range, along with a greater number of outliers, indicating more heterogeneous and occasionally long-lived bridging events. As illustrative examples, we analyzed the contact lifetimes of Asp–Na^+^–Asp and Glu–Na^+^–Glu interaction motifs (Fig. S9D). In the Asp–Na^+^–Asp case, two Asp residues remain in close proximity and collectively recruit a Na^+^ ion, which remains associated over an extended portion of the trajectory. In contrast, for the Glu–Na^+^–Glu motif, both Glu residues initially bind the same Na^+^ ion from ∼100 to 250 ns, forming a bridging interaction. After both contacts dissociate, one Glu (blue) transiently binds the ion, followed by the second Glu (orange) from ∼400 to 800 ns. Upon dissociation, the first Glu rebinds, resulting in a sequential exchange of coordination rather than simultaneous binding. Such sequential coordination is in line with a dynamic reshuffling of contacts within condensates.^20,43^ Furthermore, we identified an ion-mediated interaction network involving two Asp residues bridged by two Na^+^ ions that remain in persistent contact throughout the trajectory (Fig S10, Movie S3). In this configuration, a third Na^+^ ion engages only transiently with an additional side-chain carboxylate oxygen. This network highlights the distinctive coordination chemistry of Asp residues, which can simultaneously stabilize interactions through both backbone and side-chain carboxylate groups, enabling multivalent and dynamic ion-mediated contacts within the condensate. Finally, we observed Cl^−^-mediated bridging is largely suppressed. No significant Lys–Lys bridging is observed, likely due to the relatively low abundance of Lys residues in the system (Fig. 8B). Arg–Arg bridging is detectable but minimal, whereas Arg–Lys pairs show sporadic bridging events reflected as outliers in the distribution.

We conclude that Na^+^ is not only more abundant but also forms more frequent and longer-lived associations with residue side chains compared to Cl^−^, highlighting its dominant role in residue-ion interactions in the MUT-16 FFR condensate.

### Water bridges mediate interactions between polar uncharged residues

We next examined the presence, coordination, and role of water in the dynamics of the MUT-16 FFR condensate. Average density profiles were computed from 10 independent replicas, with uncertainty represented by shaded regions. The relatively small errors indicate that, although the condensate is dynamic, its overall density profile remains stable over the timescales sampled.

The MUT-16 FFR condensate is highly solvated (Fig. 9A), with the water density decreasing from bulk values of approximately ∼ 1000 mg/mL to ∼ 500 mg/mL within the condensate interior. This level of hydration is consistent with previous atomistic studies of protein condensates reported by Zheng et al.^19^ In contrast, the protein density within the MUT-16 condensate (∼ 800 mg/mL) is substantially higher than that reported for FUS condensates (∼ 500 mg/mL), highlighting differences in sequence composition and the force-field framework used prior to backmapping. The water density within the condensate is comparable to the density of polar residues, which is approximately ∼ 400 mg/mL. Density profiles of positively and negatively charged residues largely overlap, indicating similar spatial distributions. Notably, charged residues are preferentially localized within the condensate interior rather than at the interface with the surrounding aqueous phase. Polar residues also exhibit reduced density at the interface but display a more uniform distribution between the interface and the condensate bulk compared to charged residues. The condensate is further enriched in basic residues, with an average density of approximately ∼ 100 mg/mL, which likely contributes to the accumulation of Na^+^ ions within the condensate compared to Cl^−^ (Fig. S7). Across acidic, basic, and polar residue classes, the density profiles exhibit a common qualitative trend: a depletion at the interface, followed by a modest increase just inside the condensate, and a gradual decrease toward the condensate core.

**Figure 9:**
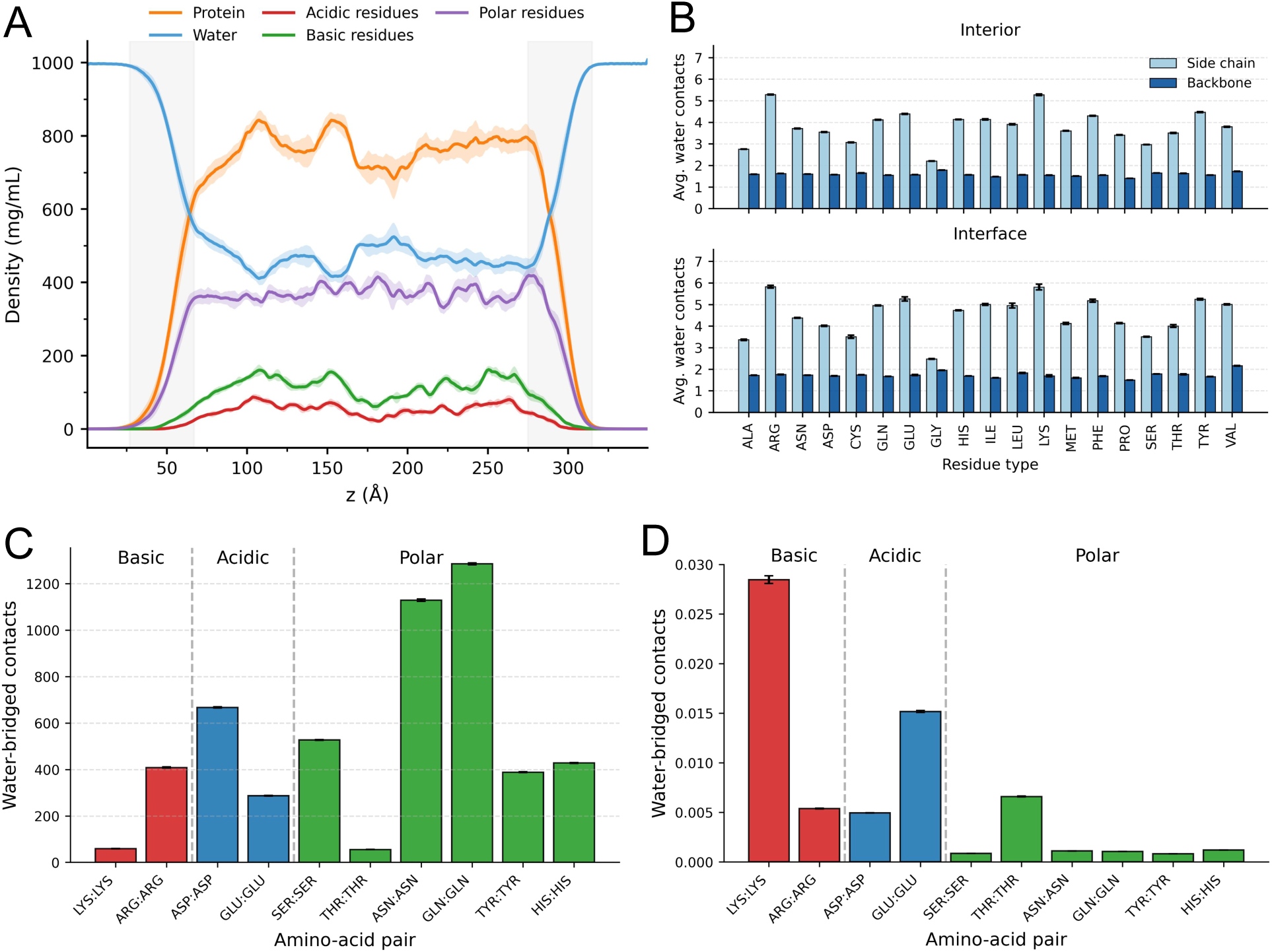
Interactions of water molecules within the MUT-16 FFR condensate. **A.** Density profiles of protein, water, and acidic, basic, and polar residues, averaged over ten independent simulation replicas. Condensate interface indicated by the shaded areas. **B.** Water–amino acid contacts in the condensate interior and at the condensate–solvent interface. **C.** Total number of water-bridged contacts among basic, acidic, and polar residue pairs. **D.** Water-bridged contact frequencies normalized by residue-pair abundance.

We further analyzed the average number of water molecules interacting with each residue, distinguishing between side-chain and backbone contributions and between the condensate interior and the condensate–solvent interface (Fig. 9B). Overall, backbone-associated water interactions are consistently lower than side-chain-associated interactions across all residue types, and backbone hydration levels are relatively uniform, as expected given the chemical similarity of peptide backbones. Backbone hydration changes minimally between the condensate interior and interface; however, all amino acid backbones are slightly more solvated at the interface, with an average increase of ∼ 0.14 water molecules compared to the interior. Among all residues, Lys and Arg exhibit the highest degree of side chain hydration, interacting on average with approximately five water molecules per residue in the condensate interior and about six water molecules at the interface. These interactions are followed in magnitude by Glu, Gln, and Tyr. Notably, several aromatic and aliphatic residues, including Phe, Val, Ile, and Leu, also display a moderate level of water association. Collectively, the average number of water contacts strongly correlates with side-chain volume in both the condensate interior (*r* = 0.88) and at the interface (*r* = 0.90) (Fig. S11). Notably, Lys and Arg exhibit higher hydration levels than expected based on this correlation, whereas Met and Gln are less hydrated than predicted by their side-chain volume, particularly within the condensate interior (Fig. S11). These observations suggest that residue hydration within the condensate is influenced not only by side-chain polarity but also by side-chain size and solvent accessibility, with larger side chains presenting greater surface area for transient water interactions.

We next analyzed the role of water in mediating bridging interactions between similarly charged residues, focusing on basic (Lys–Lys and Arg–Arg) and acidic (Asp–Asp and Glu–Glu) residue pairs that are otherwise unlikely to interact directly due to electrostatic repulsion. Both unnormalized and abundance-normalized water-bridged contact frequencies were evaluated (Fig. 9C,D). Among basic residue pairs, unnormalized water-mediated Arg–Arg interactions are more frequent than Lys–Lys interactions (Fig. 9C), which is consistent with the higher abundance of Arg residues in the system. After normalization by residue-pair abundance, however, Lys–Lys interactions exceed Arg–Arg interactions (Fig. 9D). Given that Lys–Lys interactions are rarely bridged by Cl^−^ ions (Fig. 8C), water molecules likely play a dominant role in mediating interactions between Lys residues. Such close proximity of two positive charge residues is likely made possible, at least in part, by the many negatively charged groups in the condensate (Fig. 9A). For acidic residue pairs, unnormalized Asp–Asp interactions are more frequently water-bridged than Glu–Glu interactions, reflecting the approximately 2.5-fold higher abundance of Asp relative to Glu in the sequence. In contrast, normalized contact frequencies reveal that Glu–Glu interactions are approximately threefold more likely to be bridged by water than Asp–Asp interactions, indicating a stronger intrinsic propensity of Glu residues to engage in water-mediated interactions.

We further examined water-mediated bridging interactions among polar amino acids, including Ser, Thr, Asn, Gln, Tyr, and His, which can interact directly but may also associate via intervening water molecules (Fig. 8C,D). In the unnormalized analysis, water-bridged interactions are most frequent for Asn–Asn and Gln–Gln pairs, consistent with the high abundance of these residues in the system. These are followed by Ser–Ser, His–His, and Tyr–Tyr interactions, while Thr–Thr interactions are least frequent. After normalization by residue-pair abundance, however, Thr–Thr interactions emerge as the most prominent water-bridged contacts (Fig. 8D), despite Thr exhibiting lower average hydration compared to Gln, Asn, Tyr, and His (Fig. 8B). This observation suggests that Thr residues possess a higher intrinsic propensity for forming water-mediated interactions, likely reflecting favorable geometric arrangement of backbone and side-chain polar groups rather than overall hydration alone.

## Discussion

In this study, we have investigated the dynamics of contacts within the phase-separated MUT-16 condensate at atomic resolution. Important experimental advances have revealed key drivers of protein phase separation,^2,3,5,21,80,87^ and high-resolution experiments have begun to uncover the dynamics of interactions within condensates at the molecular scale.^17,18,20,22,43^ Atomistic molecular dynamics simulations have been widely used to dissect the molecular drivers and non-covalent interactions underlying protein phase separation.^8,19,22,26,29,38^ Atomistic studies have highlighted different non-covalent interactions,^19,26,38^ as well as the critical role of interaction lifetimes in condensate dynamics.^20,43^ To better understand the dynamics of protein condensates in a biologically relevant context, we investigated MUT-16 condensates, which serve as essential scaffolds of *Mutator foci* in *C. elegans*, where they are required for small RNA processing and transposon repression.^54^ With additional experiments, we fur-ther linked phase behavior *in vivo* and *in vitro*. Using atomistic molecular dynamics simulations, we elucidated interaction propensities and the dynamics of non-covalent interactions, and also examined the interactions of ions and water in phase-separated condensates.

We observed that the most abundant contacts are relatively weak and short-lived, with median lifetimes on the nanosecond timescale, centred around a median of 9.8 ns. Galvanetto *et al.* recently linked the fast dynamics of non-covalent interactions, and what they term dynamic re-shuffling of contacts, to the mesoscopic properties of condensates.^43^ Our results show similarly fast dynamics and are consistent with their findings and conclusions, despite slight methodological differences. The mean values in our analysis are likely somewhat overestimated, as we did not resolve extremely short-lived events (≤ 100 ps) and instead considered trajectory-spanning contacts in the statistics, unlike Galvanetto.^20,43^

We provide a novel and comprehensive characterization of the contact dynamics of different amino acids within a condensate. In general, contacts may be frequent and contribute to condensation, yet appear weak when normalized for residue pair abundance.^38^ Conversely, contacts that are prominent in normalized contact maps, although potentially strong, may be rare due to low residue abundance. Overall, we find that the lifetimes of residue-residue contacts correlates with the normalized interaction propensity, but not the unnormalized interaction propensity (Fig S2). For instance Gln and Asn contacts are very prominent in absolute terms, but these contacts have persistence times on the nanosecond time scale. By contrast Tyr-Tyr interactions are less common in absolute terms, but frequent when accounting for the numbers of Tyr residues. These interactions are also much longer lived, with mean and median life times of ∼ 65 ns and ∼ 10 ns. Interestingly, some Tyr-Tyr contacts stay close for almost the entire duration of the 1 *µ*s molecular dynamics trajectories (Fig. 5C). Within this trajectory segment, contact samples conformations are typical for a *π*-*π* interaction. These contacts are relatively short-lived compared to the long times residues stay in contact (Fig S6). In this context, it is interesting to note that *π*-*π* interactions are sometimes described very accurately by molecular dynamics simulations^34^ and thus the rapid switching of the ring conformation may reflect the true dynamics of such interactions.

Our molecular dynamics simulations also provide insight into weak contacts, which play an important role in protein phase separation,^22,87^ as well as into protein–water interactions within condensates.^88,89^ Short-lived Asn and Gln contacts primarily involve hydrogen bonds (Fig. 6). Their side chains are well solvated and form a comparable number of contacts with water molecules as Asp and Glu, respectively. Water molecules are sometimes observed to bridge between Asn-Asn and Gln-Gln side chains. For instance, Thr-Thr contacts are relatively rare, both in terms of absolute contact counts and when normalized by residue abundance. Thr is comparatively highly solvated, and Thr-Thr interactions are often mediated by bridging water molecules. Interestingly, Thr-Thr is the polar residue pair that exhibits the highest number of such bridging water molecules upon abundance-normalisation(Fig. 9D). Overall, the number of water contacts a side chain forms in the condensate depends strongly on its volume, suggesting that side chains do not remain in a single fixed conformation for extended periods (Fig. S11), neither in the condensate interior nor at the interface. In this context, it is noteworthy that Schäfer and colleagues recently showed that side-chain dynamics are not significantly slowed in condensates.^30^

Unlike FUS LCD condensates, where polar and aromatic contacts dominate, MUT-16 FFR exhibits additional interaction modes. Pro is enriched in the sequence compared to FUS LCD, and in our simulations, Pro interacts with Tyr and Phe, with these interactions being relatively long-lived, showing mean lifetimes on the 50-70 ns timescale. Tyr-Pro interactions have been identified as important contacts in protein condensates.^79^ Extensive salt-bridges between Asp and Glu with Arg and Lys, as well as with counterions, distinguish MUT-16 FFR from FUS LCD. This behavior is consistent with prior simulations of the disordered N-terminal RGG domain of LAF-1.^19^ Interactions between residue side chains and ions have also been identified in condensates of highly charged proteins.^20^ In MUT-16 FFR, most ion contacts are transient, although a subset persists for hundreds of nanoseconds. We also observe dynamic reshuffling, where a Glu-Na^+^ contact is replaced by a new Glu-Na^+^ interaction via a transient bridging state (Fig. S9D). Interestingly, Na^+^ binds both Asp side chains and the backbone, and can bridge side-chain and backbone carbonyl groups of Asn and Thr, stabilizing these contacts via electrostatic coordination.

*In vitro* phase separation experiments show that MUT-16 FFR condensates dissolve at high temperatures and follow UCST behavior (Fig. 3). This observation further suggests MUT-16 phase separation underpins the formation of *Mutator foci* in *C. elegans*, considering that *Mutator foci* dissolve at high temperature. Because MUT-16 fails to phase separate without FFR, the cation–*π*, *π* stacking, and salt-bridge interactions that drive MUT-16 FFR condensation likely also govern full length MUT-16 condensate formation *in vivo* ^51^ and *in vitro*.^39^ In full length MUT-16, charged contacts are likely even more important. Residues 633-772 are enriched in Arg and Lys and mediate recruitment of MUT-8 and consequently MUT-7, an exoribonuclease essential for 22G RNA processing.^39,54^ Since, the net charge per residue (NCPR) is similar in MUT-16 full length (−0.022) and MUT-16 FFR (−0.023),^90^ we expect that Na^+^ ions to bridge between negatively charged groups of full-length MUT-16 too. Finally, the approach outlined here enables exploration of whether relationships between contact prevalence, propensity, and lifetimes generalize across diverse sequences, including human IDRs. To overcome the computational cost of residue-resolved analyses, we implemented a parallelized, unified pipeline for efficient and reproducible quantification of contact frequencies and lifetimes.

## Data and Software Availability

Scripts and data related to this publication are available at https://github.com/comp-mol-biol/cascade_computing/tree/MUT16_FFR_v0.1 and

https://doi.org/10.5281/zenodo.19219063, respectively.

## Acknowledgement

This project was funded by SFB 1551 Project No. 464588647 of the DFG (Deutsche Forschungsgemeinschaft). L.S.S. acknowledges support by ReALity (Resilience, Adaptation and Longevity), M^3^ODEL and Forschungsinitiative des Landes Rheinland-Pfalz. The authors gratefully acknowledge the data storage facilities provided by the Institute for Quantitative and Computational Biosciences (IQCB) at Johannes Gutenberg University Mainz. This work was supported by the Max Planck Graduate Center with the Johannes Gutenberg-Universität Mainz (MPGC) We gratefully acknowledge the advisory services offered and the computing time granted on the supercomputers Mogon II at Johannes Gutenberg University Mainz, which is a member of the AHRP (Alliance for High-Performance Computing in Rhineland Palatinate) and the Gauss Alliance e.V. L.S.S. thanks, Dr R. Sprangers and Dr. S. M. Hanson, for inspiring discussions. We thank Dr. V. Bussetto and Dr. S. Falk for providing the MUT-16 FFR sample.

## Human IDRs Similar to MUT-16 FFR Based on EBA_min_ and Sequence Composition

Human IDRs were ranked by EBA_min_ and RMSE to identify sequences most similar to the MUT-16 FFR. EIF2D_HUMAN (UniProt: P41214) ranked highest by EBA_min_, while CDK19_HUMAN (UniProt: Q9BWU1) was the closest match by RMSE (Fig. S1). Both IDRs lack Trp, consistent with MUT-16 FFR and FUS LCD. Whereas EIF2D_HUMAN de-viates in the abundance of several residues (notably Ile, Lys, Leu, and Glu), CDK19_HUMAN closely matches the overall sequence composition of MUT-16 FFR, including similar levels of charged residues (Arg, Lys, Glu, and Asp).

## Pair-specific analysis of Glu-Arg and Asp-Arg contacts

To better understand how the dynamics of Glu-Arg and Asp-Arg contacts, and whether Asp-Arg interactions really tend to be longer lived than Glu-Arg interactions (Fig S2D), we have looked at specific residue pairs. This gives an even more detailed view of the contact dynamics than the grouping of contacts by pair-wise residue types (Eq. 1). In this residue-specific analysis, we still group together contacts across different copies of the protein chain, averaging across the 65 copies of MUT-16 FFR in our simulations, and we average across the ten simulation trajectories. In the analysis of the contacts and the dynamics of IDRs one often averages across protein chains to understand contacts^19,38^ and dynamics^18,43^ on per-residue basis. For instance, Galvanetto have computed per-residue mean-square displacements.^43^

Overall, we see hundreds of events even for specific Arg-Glu and Arg-Asp pairs (Fig.**9.** D,G), with just a single Glu-Arg pair for which we observed less than a hundred events. Looking at the individual Glu-Arg and pairs we see that contacts are have median lifetimes ranging from 9.3 ns to 28.2 ns and mean lifetimes from 43.5 ns to 123.5 ns. The Asp-Arg interactions are longer lived with median lifetimes 10.8 ns to 45.0 ns and mean lifetimes 54.3 ns to 197.3 ns. For the longest-lived Glu-Arg pair, Glu 869-Arg 875, we observed 240 contact events, with a median and mean lifetime of 28.2 and 123.5 ns, respectively. For the longest-lived Asp-Arg pair, Asp 917-Arg 904, we detected 121 contact events, with median and mean lifetime of 45.0 ns and 197.3 ns, respectively. For this Asp-Arg pair, approximately 15% of the events have durations of almost 800 ns. The long-lived Glu-Arg pair: the fraction of events that are almost trajectory-spanning was closer to 10%. The contact statistics of these two pairs are consistent with the notion that Glu-Arg pairs form more frequently than Asp-Arg pairs, due to their more extended side chains, but also dissociate more readily due to their longer side chain.

---

**Table S1:**
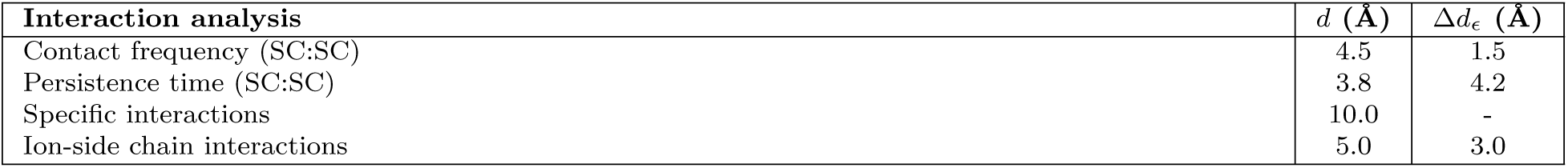
Cutoff parameters used for interaction and persistence analyses.

**Figure S1:**
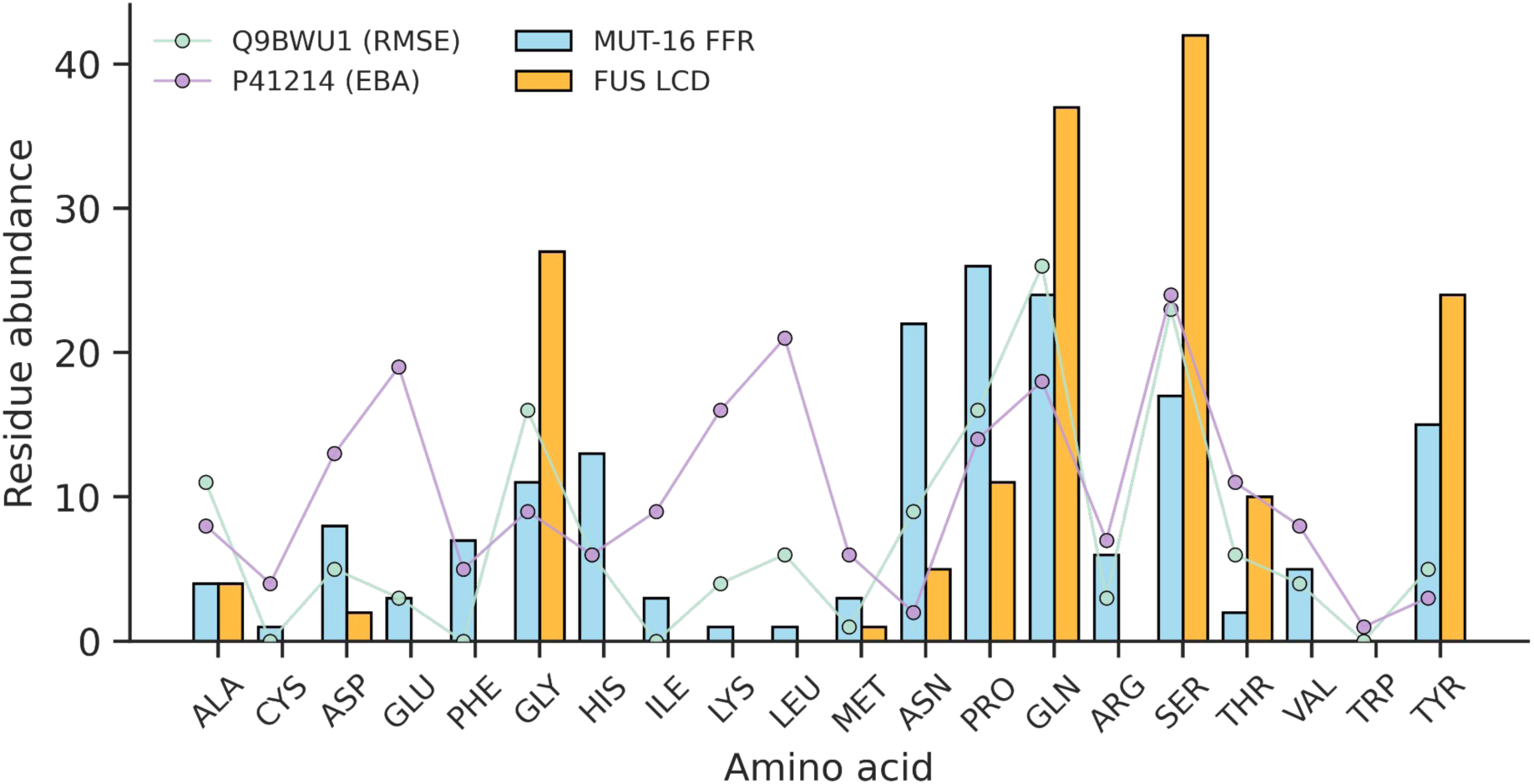
Residue abundance profiles of the MUT-16 FFR (blue) and the FUS LCD (orange). For comparison, residue abundances of representative human IDRs identified as most similar to the MUT-16 FFR based on embedding-based alignment (EBA) similarity (purple) and root-mean-square error (RMSE) (green) are overlaid as scatter points.

**Figure S2:**
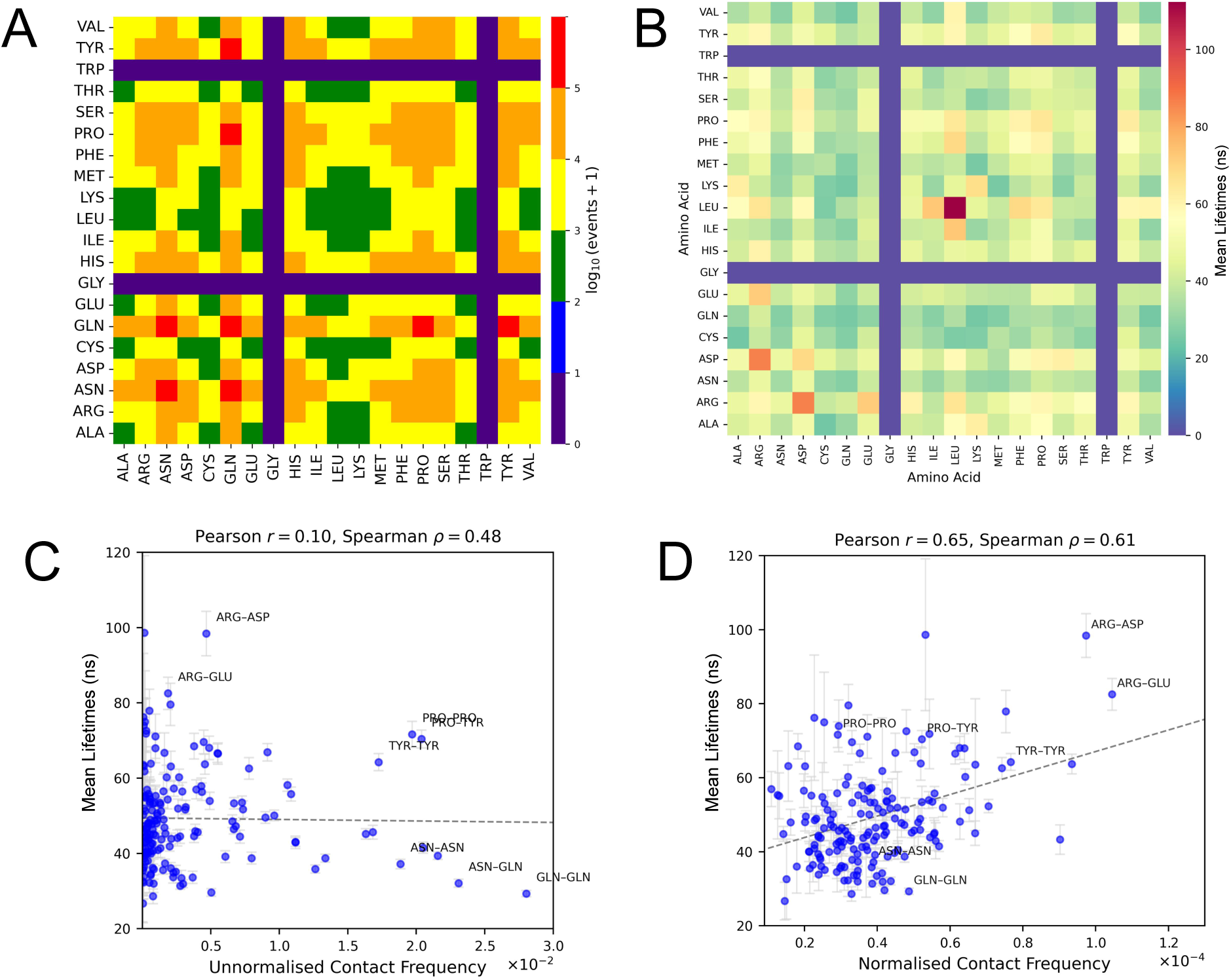
Relationship between contact frequency, persistence, and residue abundance in the MUT-16 FFR condensate. **A.** Heat map showing the total number of binding events observed for each residue-residue pair that were included in the statistical analysis of contact distributions and mean lifetimes. The color scale represents the number of events on a logarithmic scale. **B.** Heat map of the mean lifetimes of side-chain-mediated interactions, highlighting the relative stability and typical lifetimes of residue-residue contacts within the condensate. **C.** Correlation between unnormalised contact frequency and mean persistence time, illustrating how frequently formed contacts relate to their temporal stability. **D.** Correlation between abundance-normalised contact frequency and mean persistence time, isolating intrinsic interaction stability from effects arising due to residue abundance.

**Figure S3:**
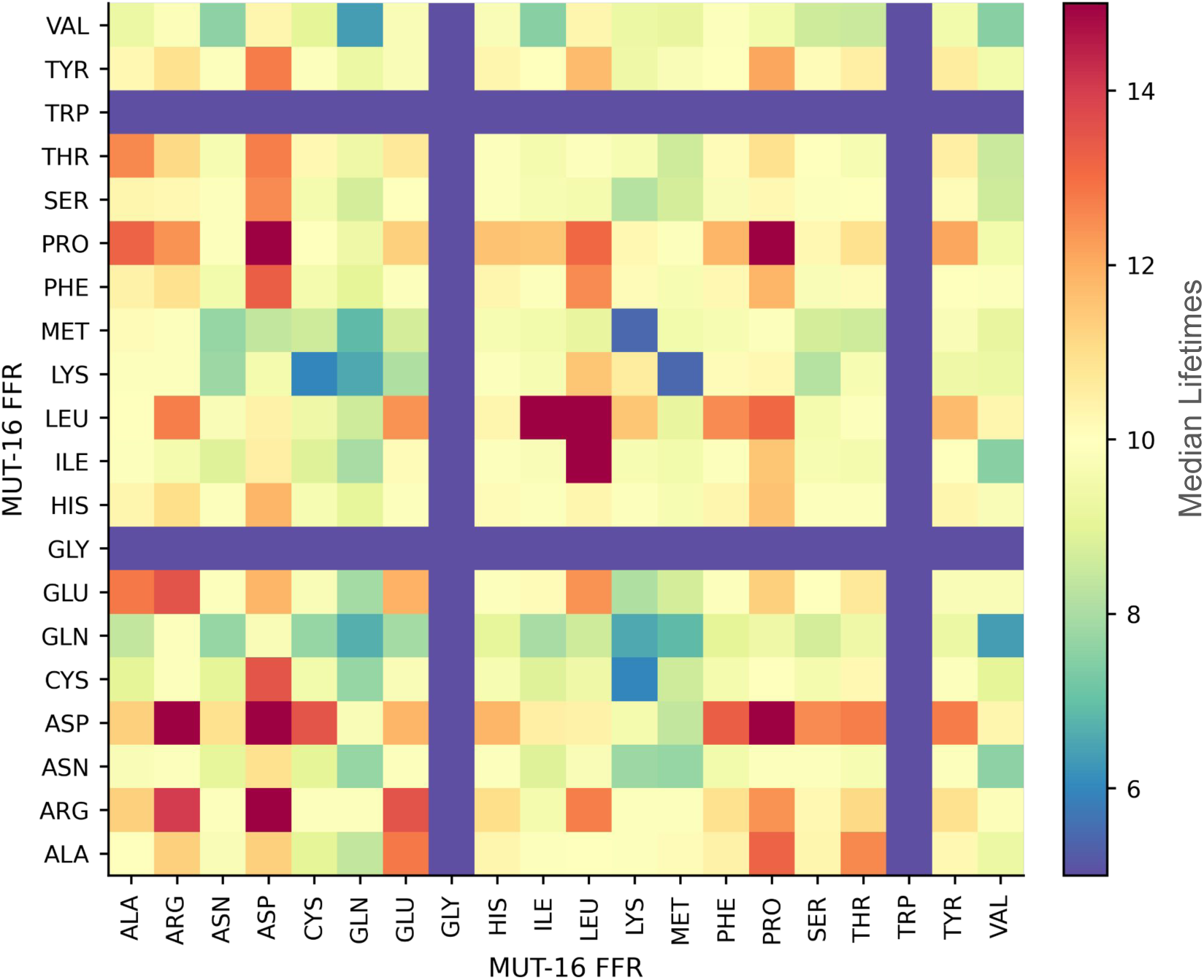
Heat map showing the median lifetimes of side-chain-mediated interactions, emphasizing the typical (central) persistence of residue–residue contacts within the condensate. By focusing on the median rather than the mean, the map captures the representative interaction timescales while minimizing the influence of rare, long-lived events, thereby providing a robust measure of relative interaction stability across residue pairs.

**Figure S4:**
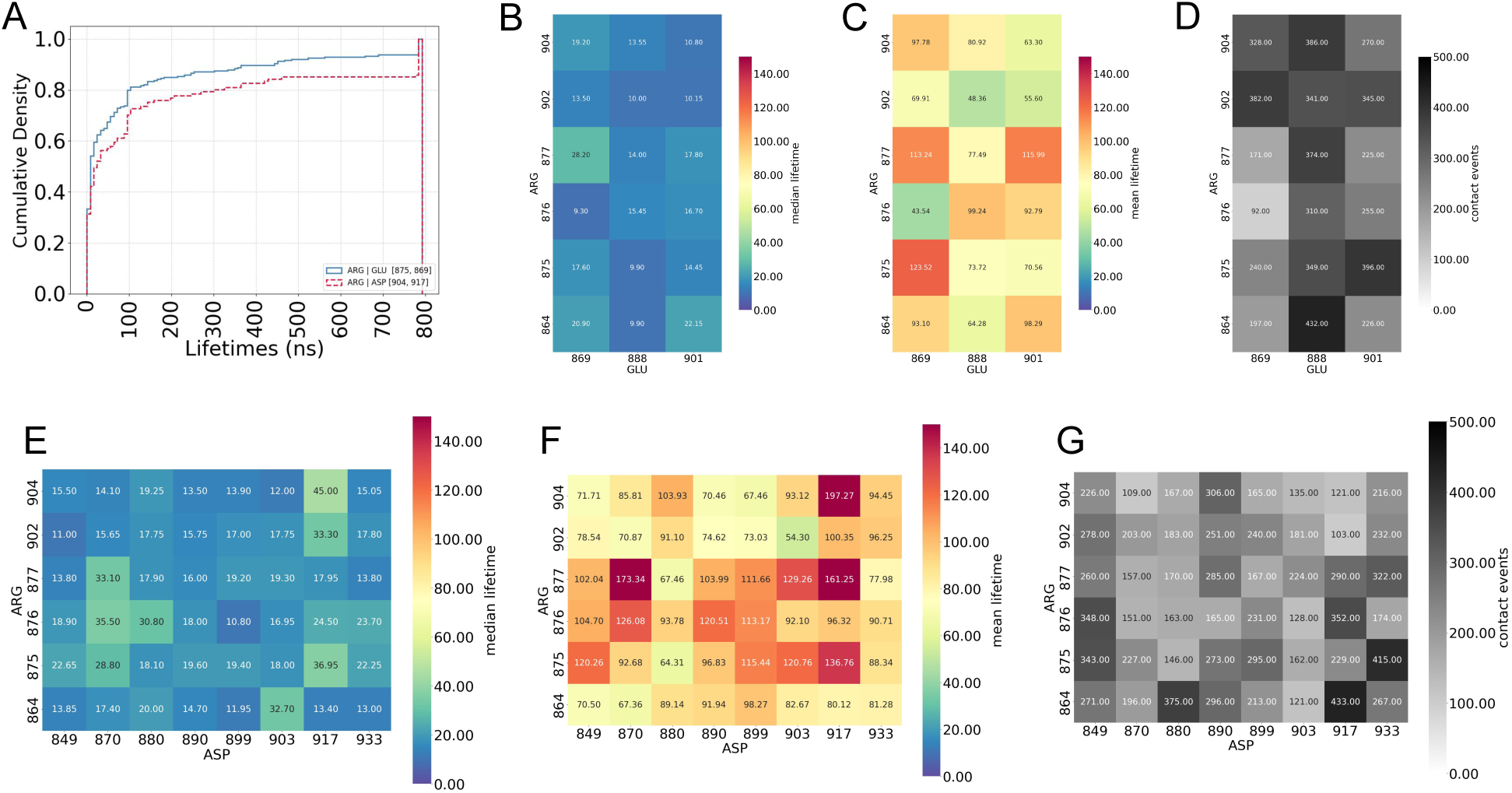
A. Cumulative lifetime distribution for Arg 875-Glu 869 and Arg 904 and Asp 917, illustrating the overall distribution of contact persistence. **B.** Median lifetimes of individual Arg-Glu residue pairs in the MUT-16 FFR sequence, highlighting the typical interaction timescales. **C.** Mean lifetimes of specific Arg-Glu residue pairs, reflecting the influence of longer-lived interactions. **D.** Number of binding events observed for Arg-Glu pairs in the MUT-16 FFR sequence, indicating interaction frequency. **E.** Median lifetimes of Arg-Asp residue pairs, providing a robust measure of typical contact stability. **F.** Mean lifetimes of specific Arg-Asp residue pairs, capturing contributions from long-lived contacts. **G.** Number of binding events for Arg-Asp pairs in the MUT-16 FFR sequence, reflecting the occurrence of these interactions across the trajectory.

**Figure S5:**
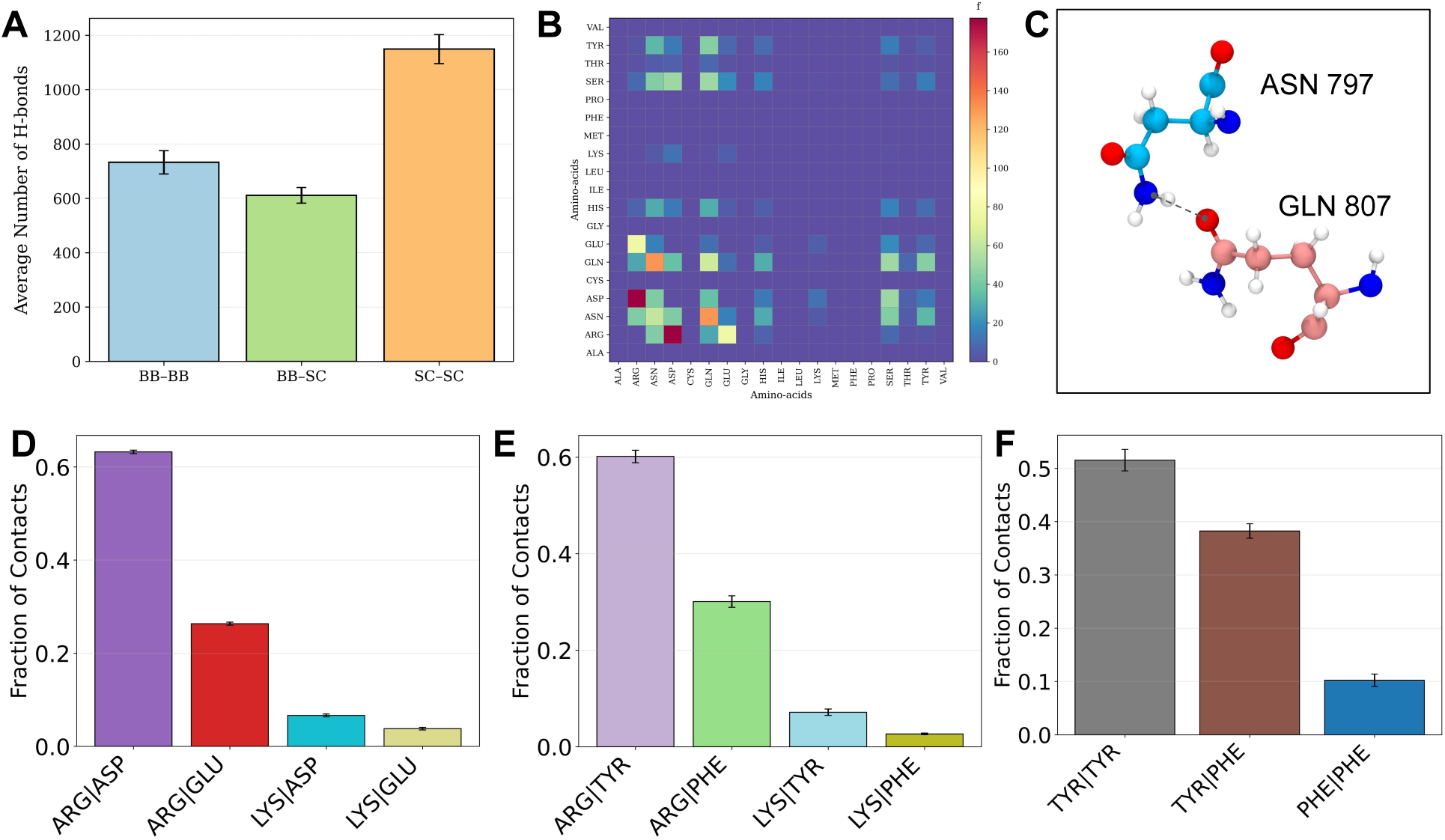
Prevalence of distinct noncovalent interaction types in the MUT-16 FFR condensate. **A.** Average number of hydrogen bonds formed between backbone-backbone (BB:BB), backbone-side chain (BB:SC), and side chain-side chain (SC:SC) contacts in the MUT-16 FFR condensate. Values are averaged over all simulation frames and across 10 independent replica trajectories, with error bars representing the standard error of the mean. **B.** Heat map of unnormalised contact frequencies for residue pairs capable of forming hydrogen bonds, reflecting the overall prevalence of hydrogen-bonded interactions. **C.** Representative snap-shot of a hydrogen bond formed between Asn and Gln residues, illustrating a typical polar side-chain interaction observed in the condensate. **D.** Bar graph showing the unnormalised fraction of contacts for residue pairs that form salt bridges, highlighting electrostatically driven interactions. **E.** Bar graph showing the unnormalised fraction of contacts for residue pairs that form cation–*π* interactions. **F.** Bar graph showing the unnormalised fraction of contacts for residue pairs that form *π*–*π* stacking interactions, emphasising the contribution of aromatic interactions to condensate organisation.

**Figure S6:**
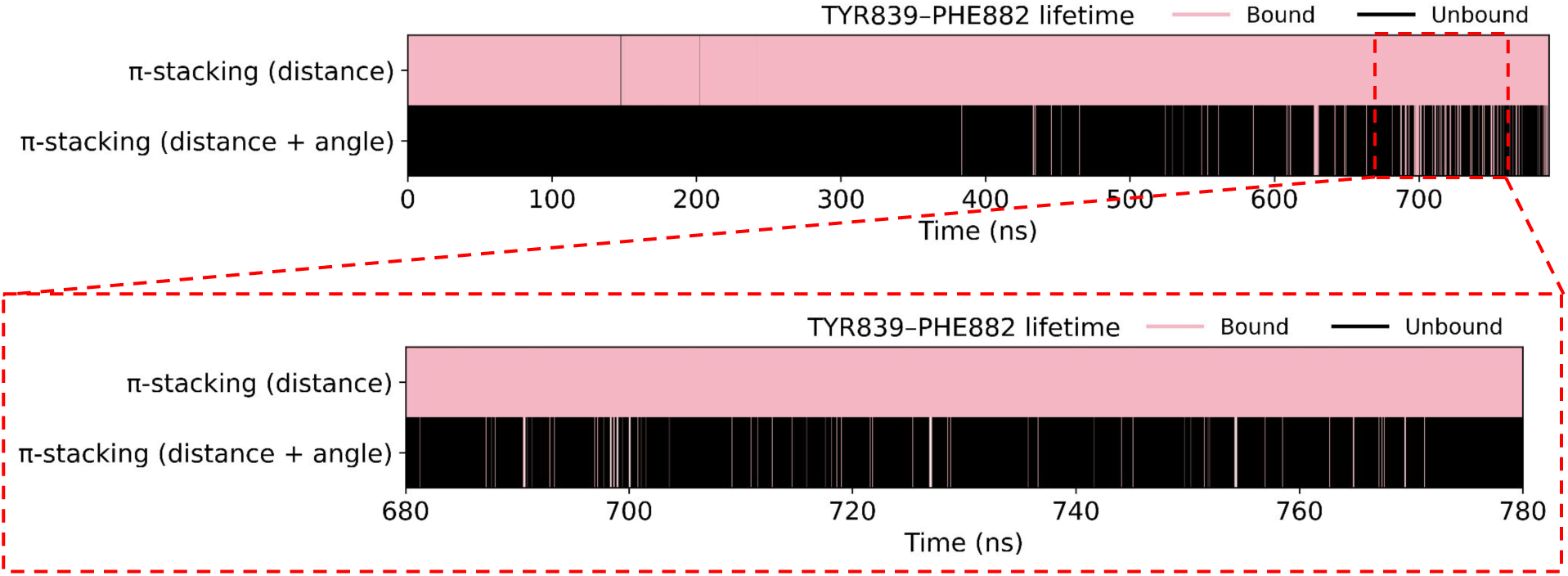
Representative lifetime of a *π*-*π* stacking interaction between Tyr and Phe residues, showing transitions between bound (pink) and unbound (black) states. Binding is identified using two criteria: a distance-only cutoff and a combined distance-and-angular cutoff, demonstrating how the inclusion of orientational constraints refines the detection of stacked configurations.

**Figure S7:**
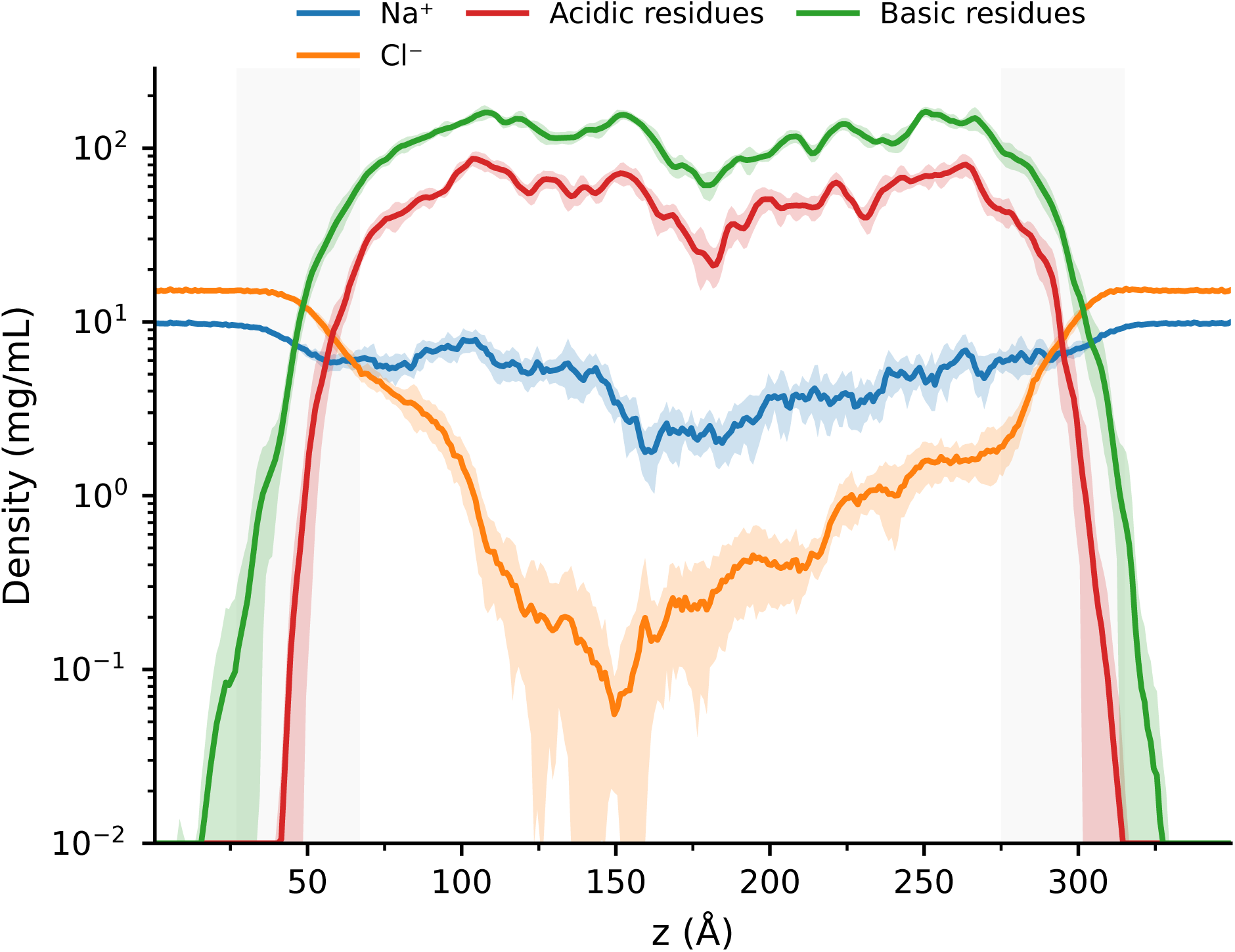
Density profiles of Na^+^ and Cl^−^ ions compared with acidic and basic residues along the condensate slab normal.

**Figure S8:**
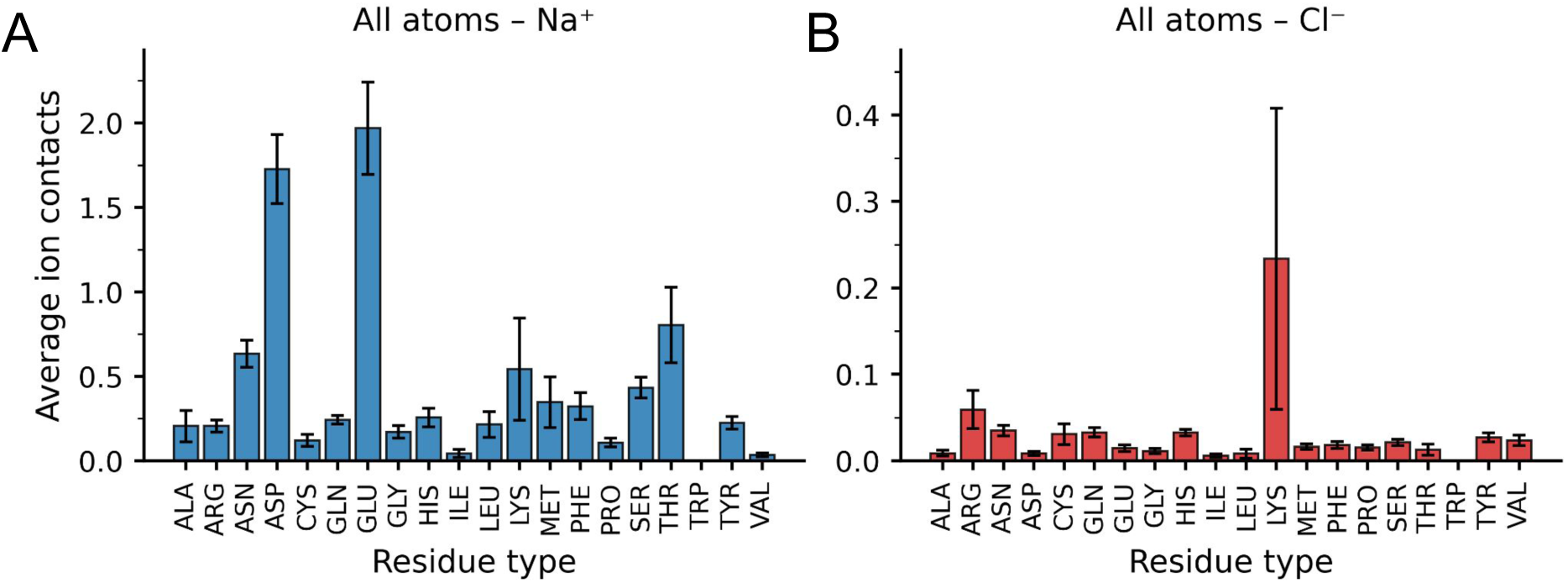
Residue-resolved ion association within the MUT-16 FFR condensate. **A.** Average number of Na^+^ ions interacting with all atoms of each residue (backbone + side chain). **B.** Average number of Cl^−^ ions interacting with all atoms of each residue (backbone + side chain).

**Figure S9:**
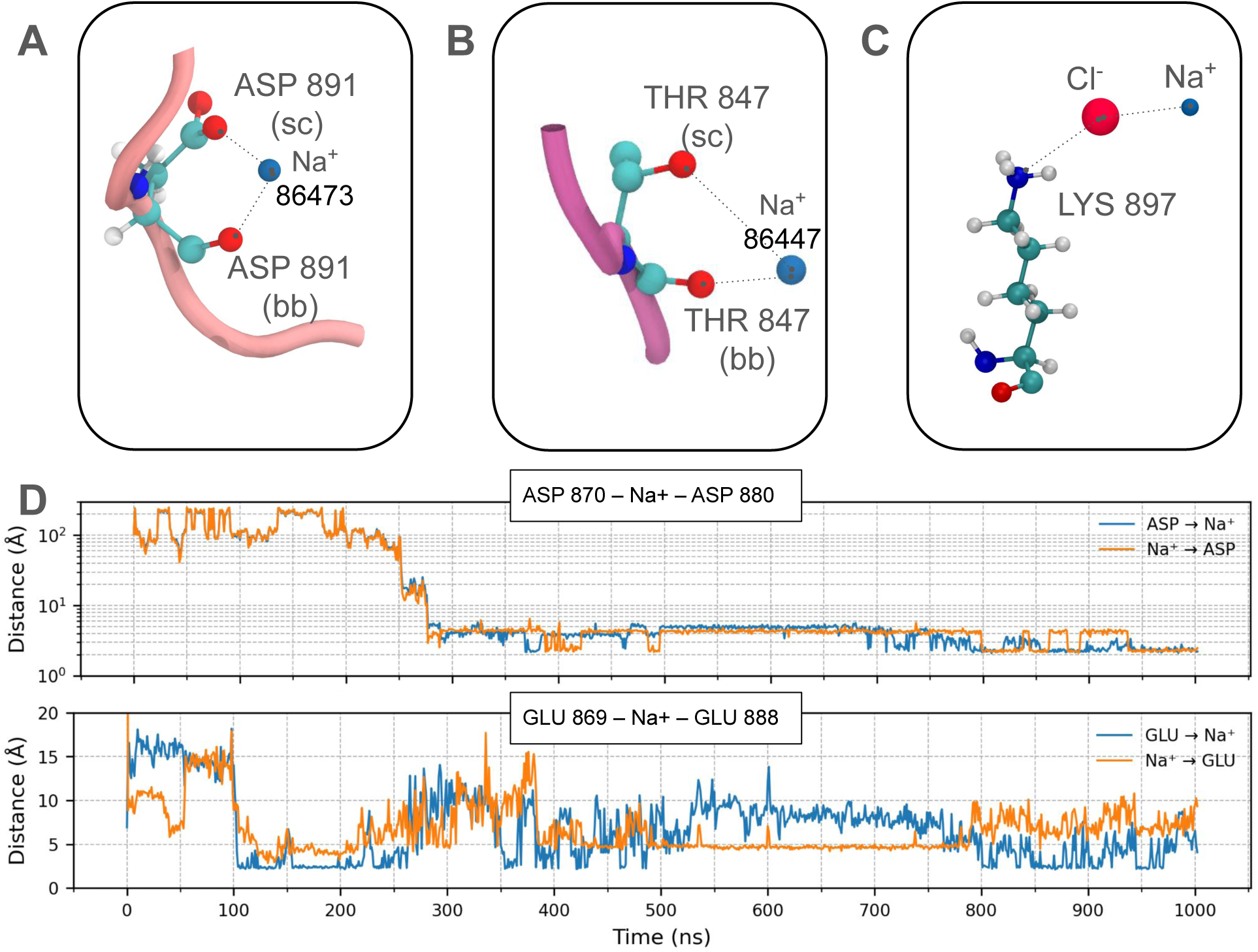
Representative ion-mediated interaction motifs observed in the MUT-16 FFR condensate. **(A)** Na^+^-mediated bridging interaction between the side chain and backbone of an Asp residue. **(B)** Na^+^-mediated bidentate interaction involving both the side chain and backbone of a Thr residue. **(C)** Interaction between Lys residues and Na^+^ facilitated by a bridging Cl^−^ ion. **(D)** Time series depicting representative Asp–Na^+^–Asp and Glu–Na^+^–Glu bridging interactions. Distances are measured between the negatively charged oxygen atoms of Asp and Glu side chains and the Na^+^ ion.

**Figure S10:**
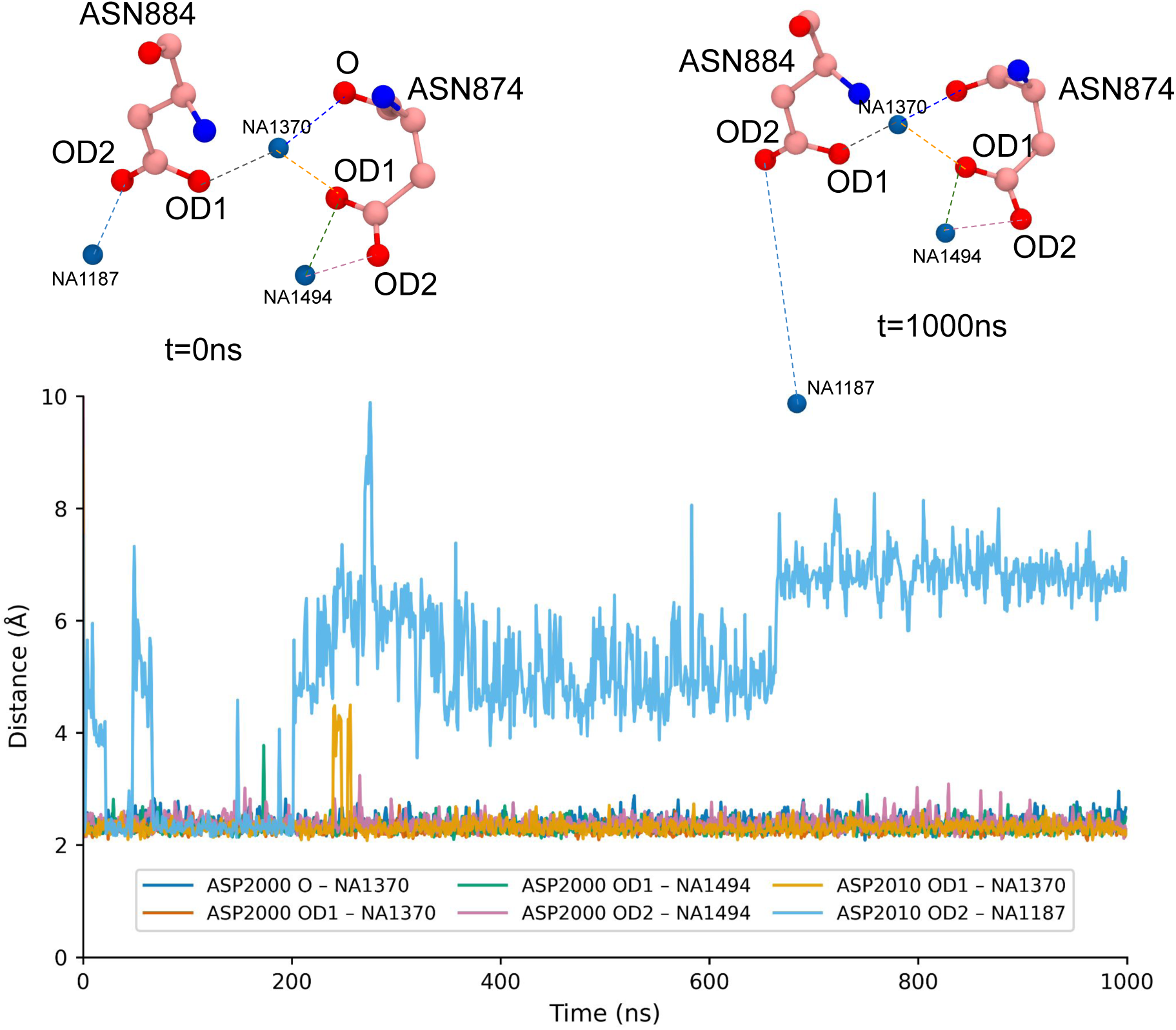
Visual representation and time series of multiple Na^+^-mediated bridging event between two Asp residues, involving both side-chain carboxylate oxygen atoms and backbone oxygen atoms. Shaded areas indicate the interface between the dense and dilute phases.

**Figure S11:**
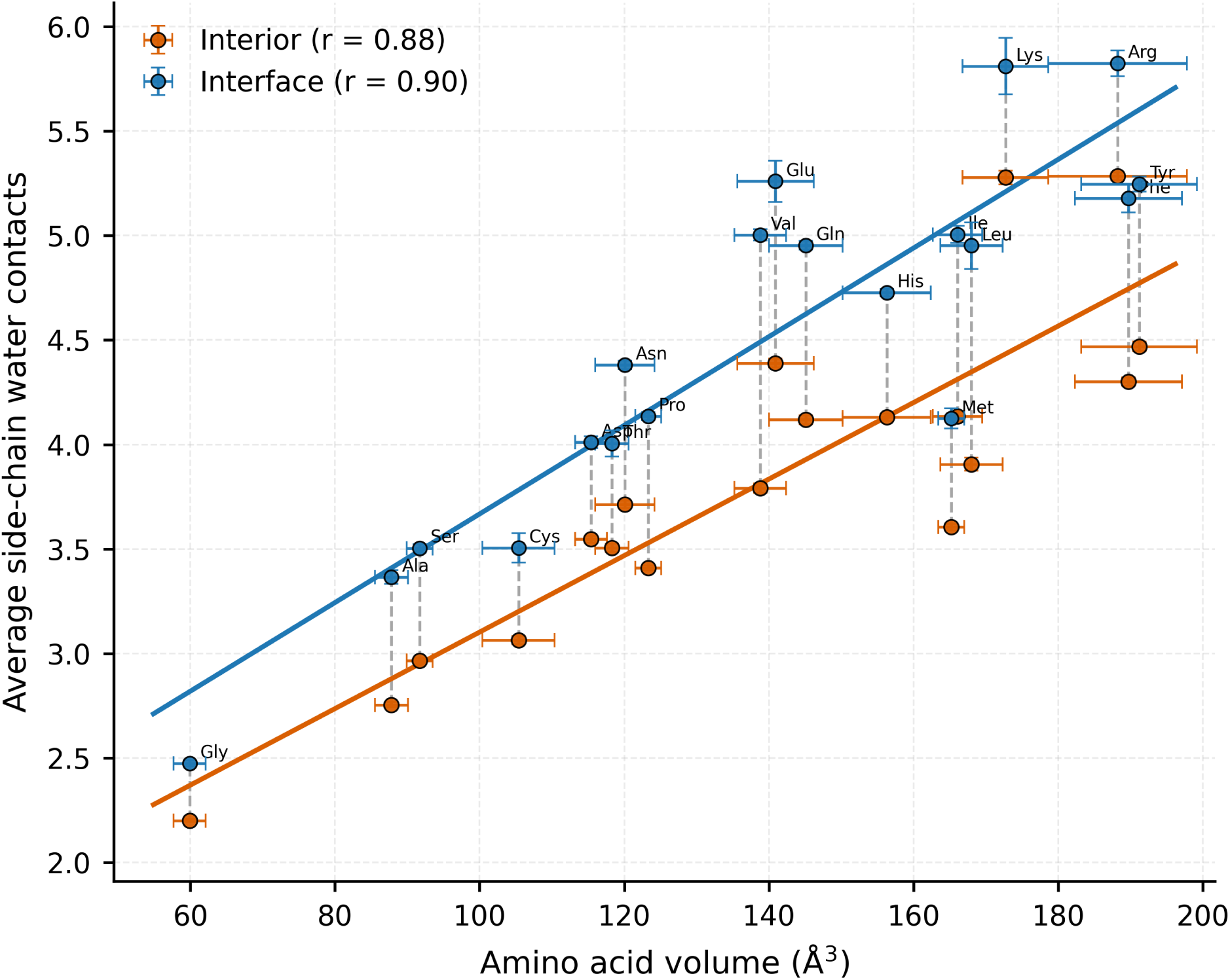
Correlation plot illustrating the relationship between the average number of water molecules interacting with amino acid side chains and the corresponding side-chain volume, evaluated separately for residues located in the condensate interior and in the bulk phase.

## Notes

https://doi.org/10.5281/zenodo.19219063

https://github.com/comp-mol-biol/cascade_computing/tree/MUT16_FFR_v0.1

### Competing Interest Statement

The authors have declared no competing interest.

### Summary of Updates

This revision has added the citation of a new pre-print, "Atomistic Simulations of Biomolecular Condensates", in ChemRxiv by Kandarp et. al.

