## Supplementary material for "Atomistic simulations reveal sub-*µ*s contact dynamics in MUT-16 condensates": Supplimentary text and figures

### Supporting information

Table S1: Cutoff parameters used for interaction and persistence analyses.

| **Interaction analysis** | $d$ (Å) | $\Delta d_{\epsilon}$ (Å) |
|---|---|---|
| Contact frequency (SC:SC) | 4.5 | 1.5 |
| Persistence time (SC:SC) | 3.8 | 4.2 |
| Specific interactions | 10.0 | - |
| Ion-side chain interactions | 5.0 | 3.0 |

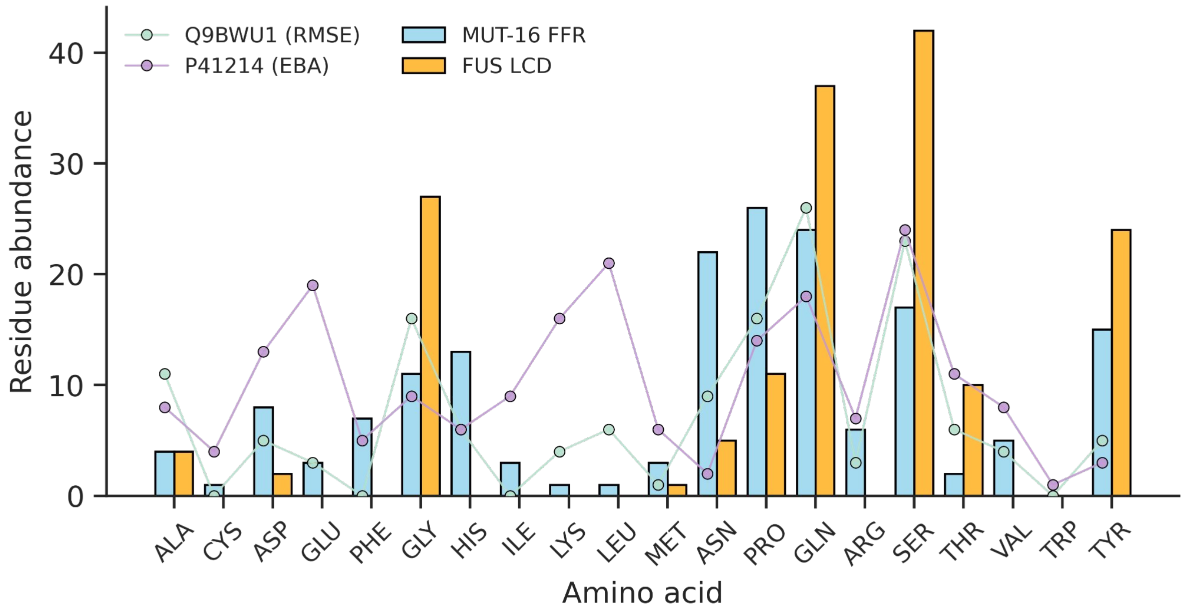


Figure S1: Residue abundance profiles of the MUT-16 FFR (blue) and the FUS LCD (orange). For comparison, residue abundances of representative human IDRs identified as most similar to the MUT-16 FFR based on embedding-based alignment (EBA) similarity (purple) and root-mean-square error (RMSE) (green) are overlaid as scatter points.

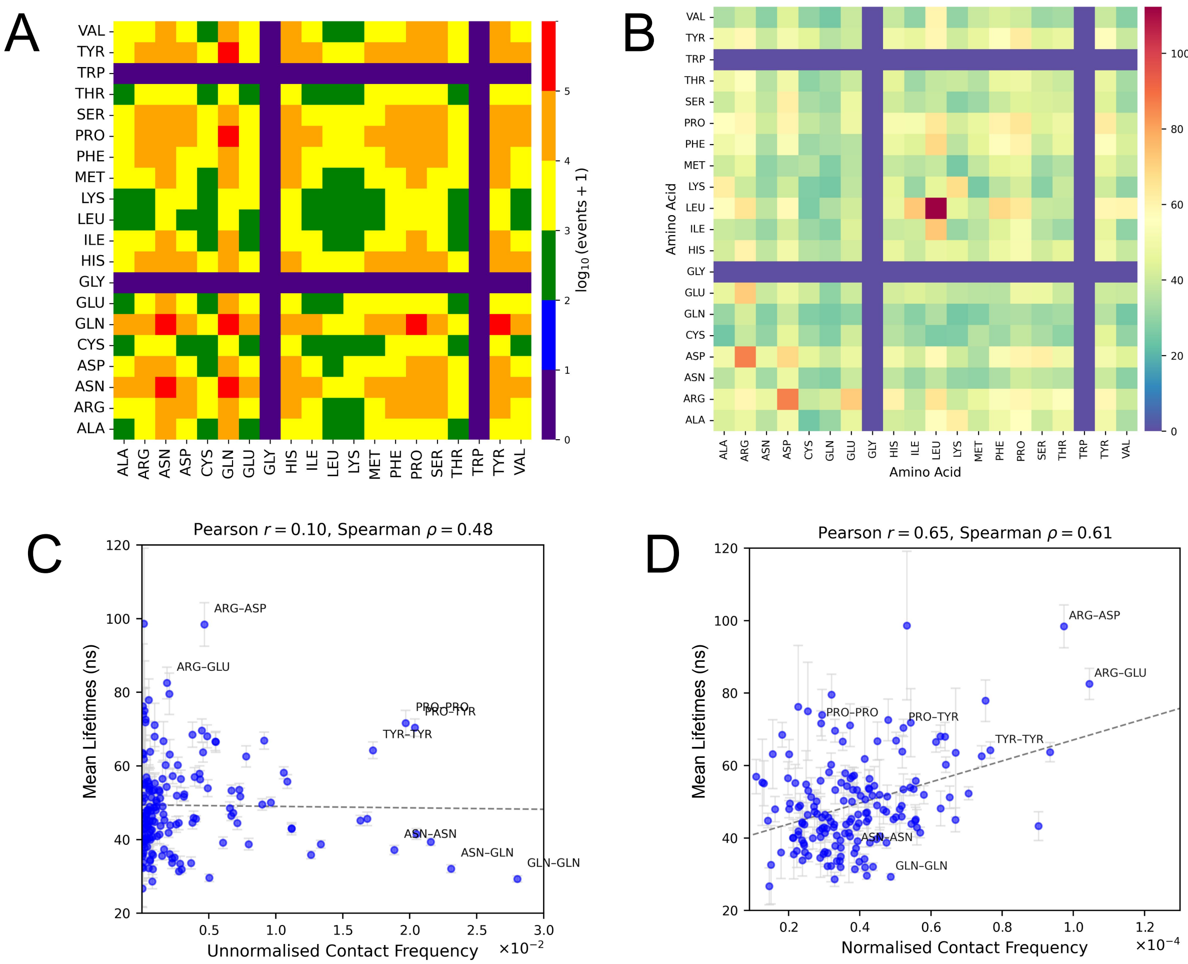


Figure S2: Relationship between contact frequency, persistence, and residue abundance in the MUT-16 FFR condensate. **A.** Heat map showing the total number of binding events observed for each residue-residue pair that were included in the statistical analysis of contact distributions and mean lifetimes. The color scale represents the number of events on a logarithmic scale. **B.** Heat map of the mean lifetimes of side-chain-mediated interactions, highlighting the relative stability and typical lifetimes of residue-residue contacts within the condensate. **C.** Correlation between unnormalised contact frequency and mean persistence time, illustrating how frequently formed contacts relate to their temporal stability. **D.** Correlation between abundance-normalised contact frequency and mean persistence time, isolating intrinsic interaction stability from effects arising due to residue abundance.

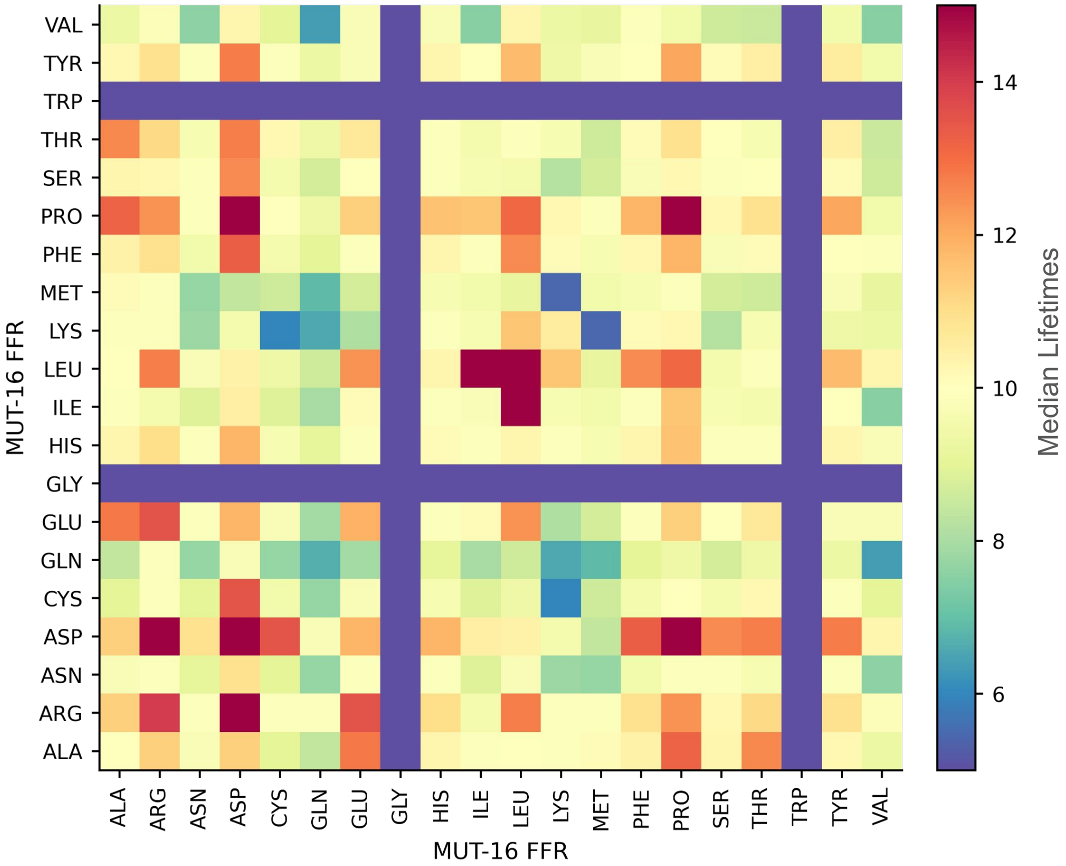


Figure S3: Heat map showing the median lifetimes of side-chain-mediated interactions, emphasizing the typical (central) persistence of residue–residue contacts within the condensate. By focusing on the median rather than the mean, the map captures the representative interaction timescales while minimizing the influence of rare, long-lived events, thereby providing a robust measure of relative interaction stability across residue pairs.

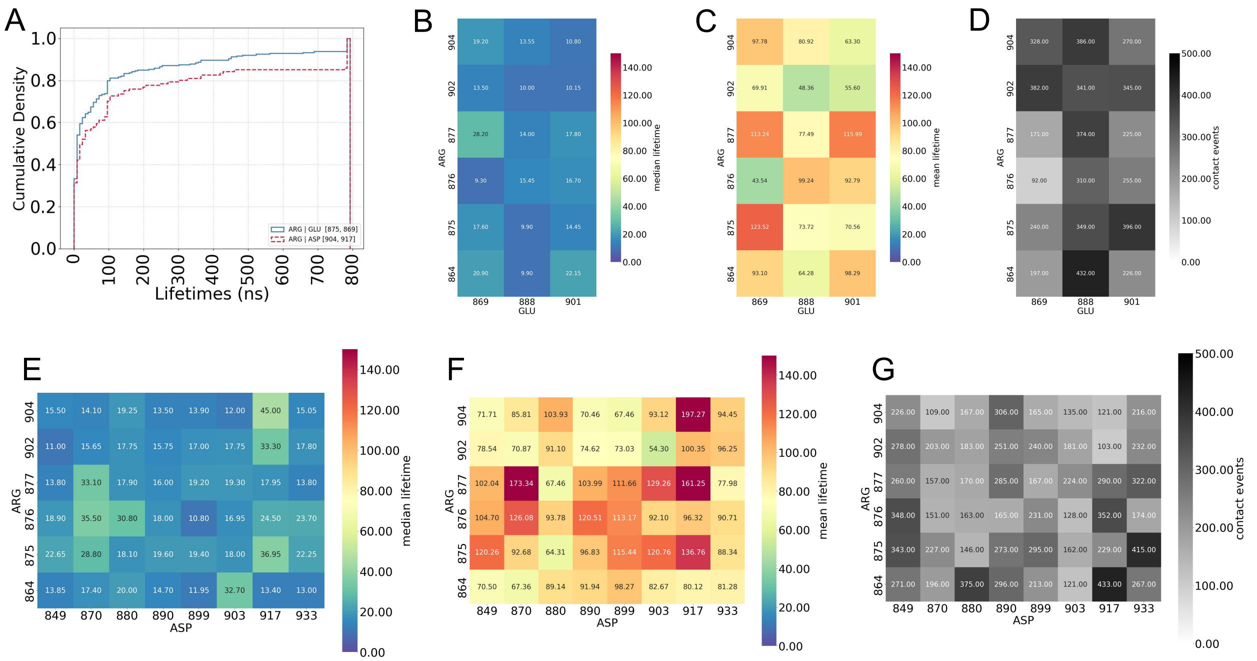


Figure S4: **A.** Cumulative lifetime distribution for Arg 875-Glu 869 and Arg 904 and Asp 917, illustrating the overall distribution of contact persistence. **B.** Median lifetimes of individual Arg-Glu residue pairs in the MUT-16 FFR sequence, highlighting the typical interaction timescales. **C.** Mean lifetimes of specific Arg-Glu residue pairs, reflecting the influence of longer-lived interactions. **D.** Number of binding events observed for Arg-Glu pairs in the MUT-16 FFR sequence, indicating interaction frequency. **E.** Median lifetimes of Arg-Asp residue pairs, providing a robust measure of typical contact stability. **F.** Mean lifetimes of specific Arg-Asp residue pairs, capturing contributions from long-lived contacts. **G.** Number of binding events for Arg-Asp pairs in the MUT-16 FFR sequence, reflecting the occurrence of these interactions across the trajectory.

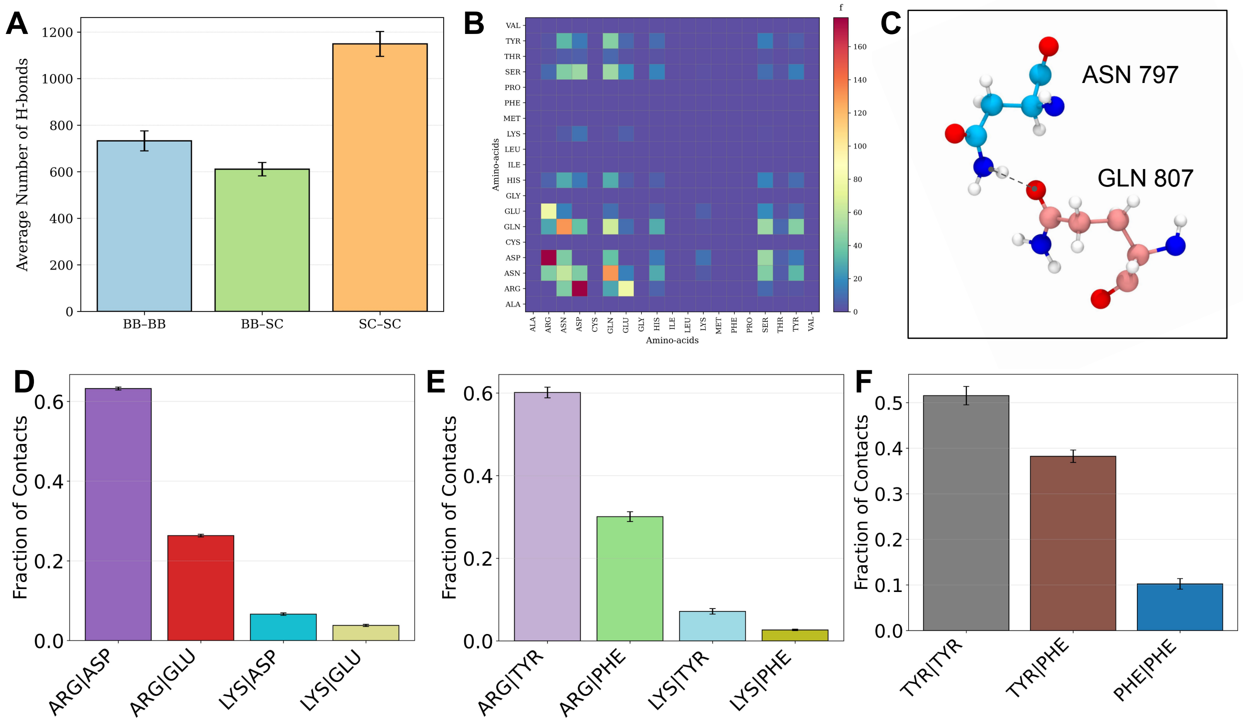


Figure S5: Prevalence of distinct noncovalent interaction types in the MUT-16 FFR condensate. **A.** Average number of hydrogen bonds formed between backbone-backbone (BB:BB), backbone-side chain (BB:SC), and side chain-side chain (SC:SC) contacts in the MUT-16 FFR condensate. Values are averaged over all simulation frames and across 10 independent replica trajectories, with error bars representing the standard error of the mean. **B.** Heat map of unnormalised contact frequencies for residue pairs capable of forming hydrogen bonds, reflecting the overall prevalence of hydrogen-bonded interactions. **C.** Representative snapshot of a hydrogen bond formed between Asn and Gln residues, illustrating a typical polar side-chain interaction observed in the condensate. **D.** Bar graph showing the unnormalised fraction of contacts for residue pairs that form salt bridges, highlighting electrostatically driven interactions. **E.** Bar graph showing the unnormalised fraction of contacts for residue pairs that form cation–$\pi$ interactions. **F.** Bar graph showing the unnormalised fraction of contacts for residue pairs that form $\pi$–$\pi$ stacking interactions, emphasising the contribution of aromatic interactions to condensate organisation.

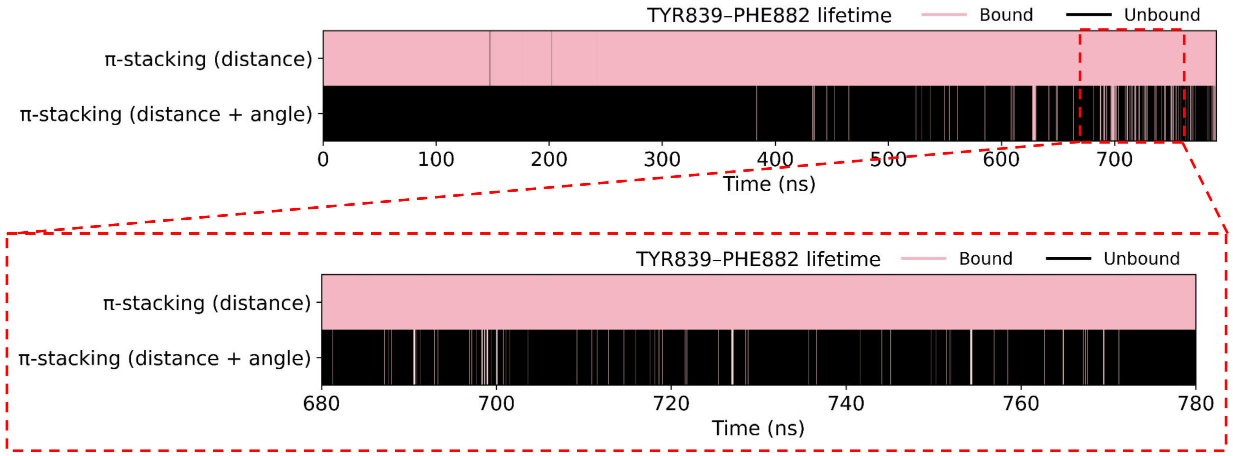


Figure S6: Representative lifetime of a $\pi$-$\pi$ stacking interaction between Tyr and Phe residues, showing transitions between bound (pink) and unbound (black) states. Binding is identified using two criteria: a distance-only cutoff and a combined distance-and-angular cutoff, demonstrating how the inclusion of orientational constraints refines the detection of stacked configurations.

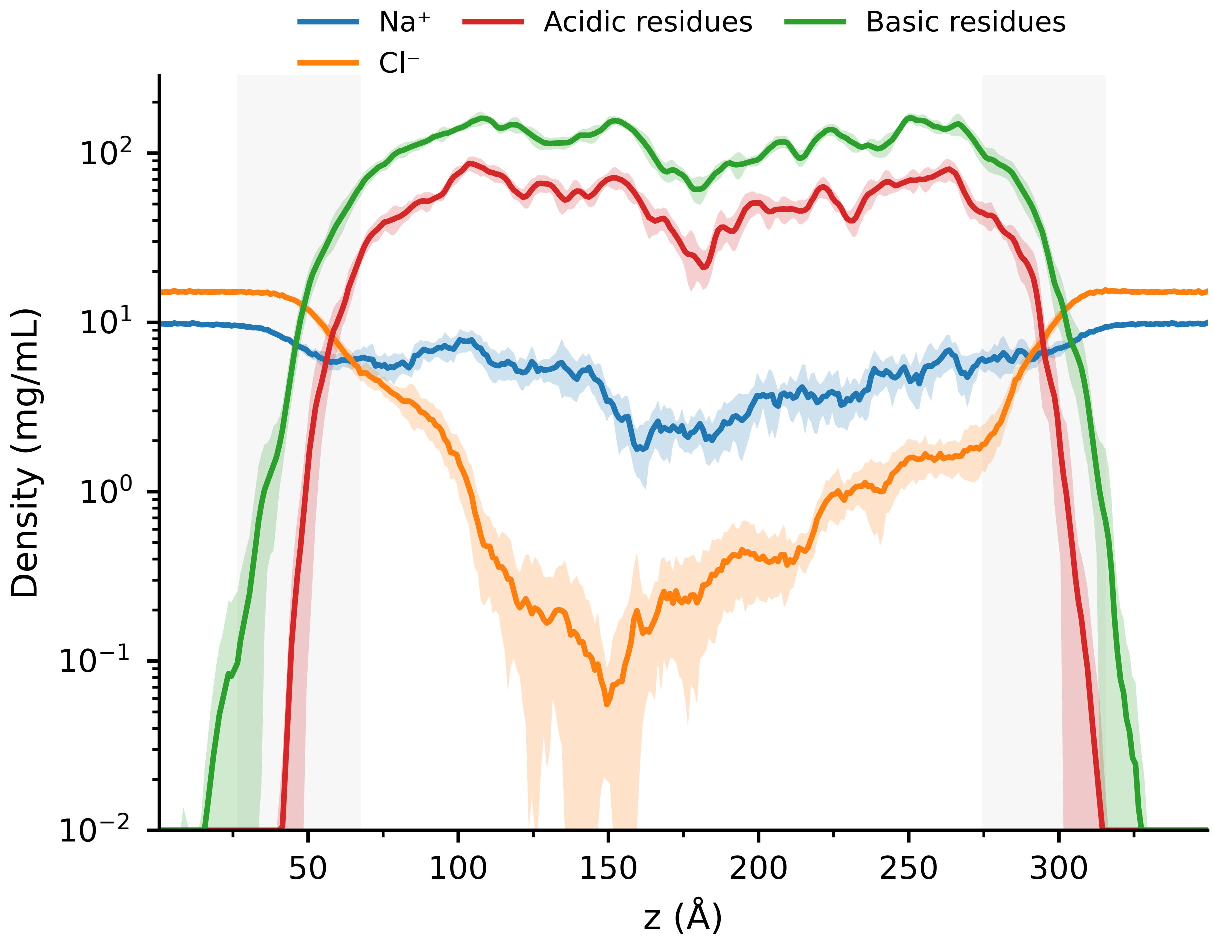


Figure S7: Density profiles of $Na^+$ and $Cl^-$ ions compared with acidic and basic residues along the condensate slab normal.

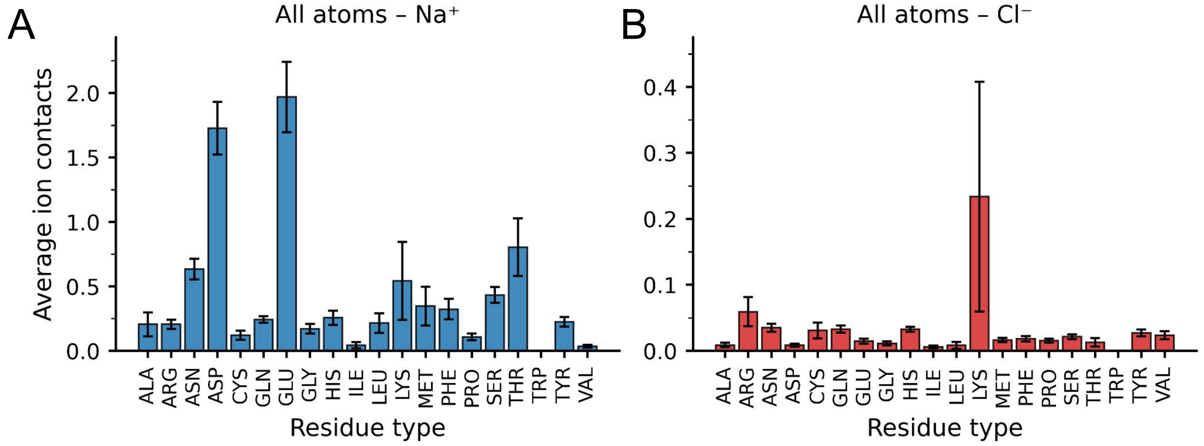


Figure S8: Residue-resolved ion association within the MUT-16 FFR condensate. **A.** Average number of $Na^+$ ions interacting with all atoms of each residue (backbone + side chain). **B.** Average number of $Cl^-$ ions interacting with all atoms of each residue (backbone + side chain).

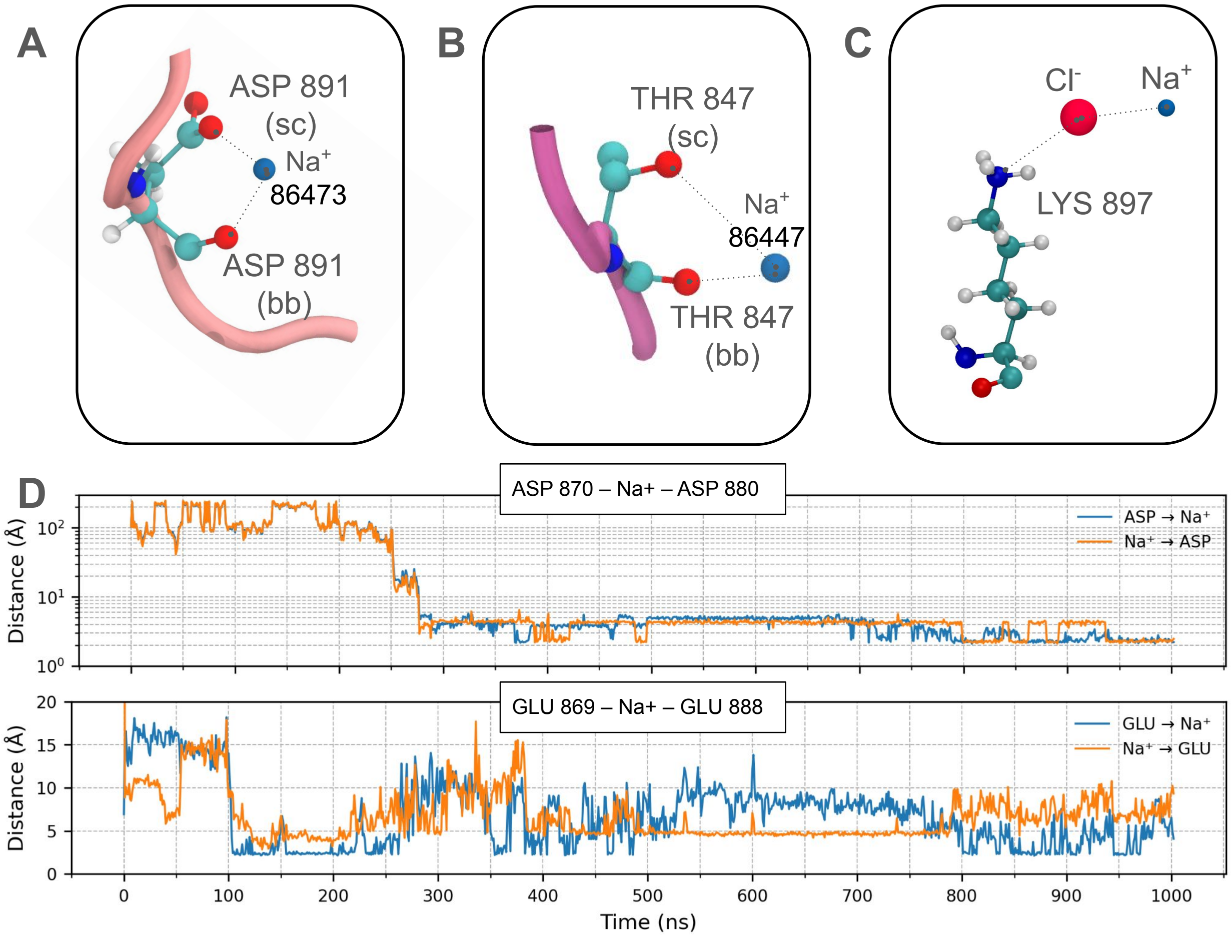


Figure S9: Representative ion-mediated interaction motifs observed in the MUT-16 FFR condensate. **(A)** $Na^+$-mediated bridging interaction between the side chain and backbone of an Asp residue. **(B)** $Na^+$-mediated bidentate interaction involving both the side chain and backbone of a Thr residue. **(C)** Interaction between Lys residues and $Na^+$ facilitated by a bridging $Cl^-$ ion. **(D)** Time series depicting representative Asp–$Na^+$–Asp and Glu–$Na^+$–Glu bridging interactions. Distances are measured between the negatively charged oxygen atoms of Asp and Glu side chains and the $Na^+$ ion.

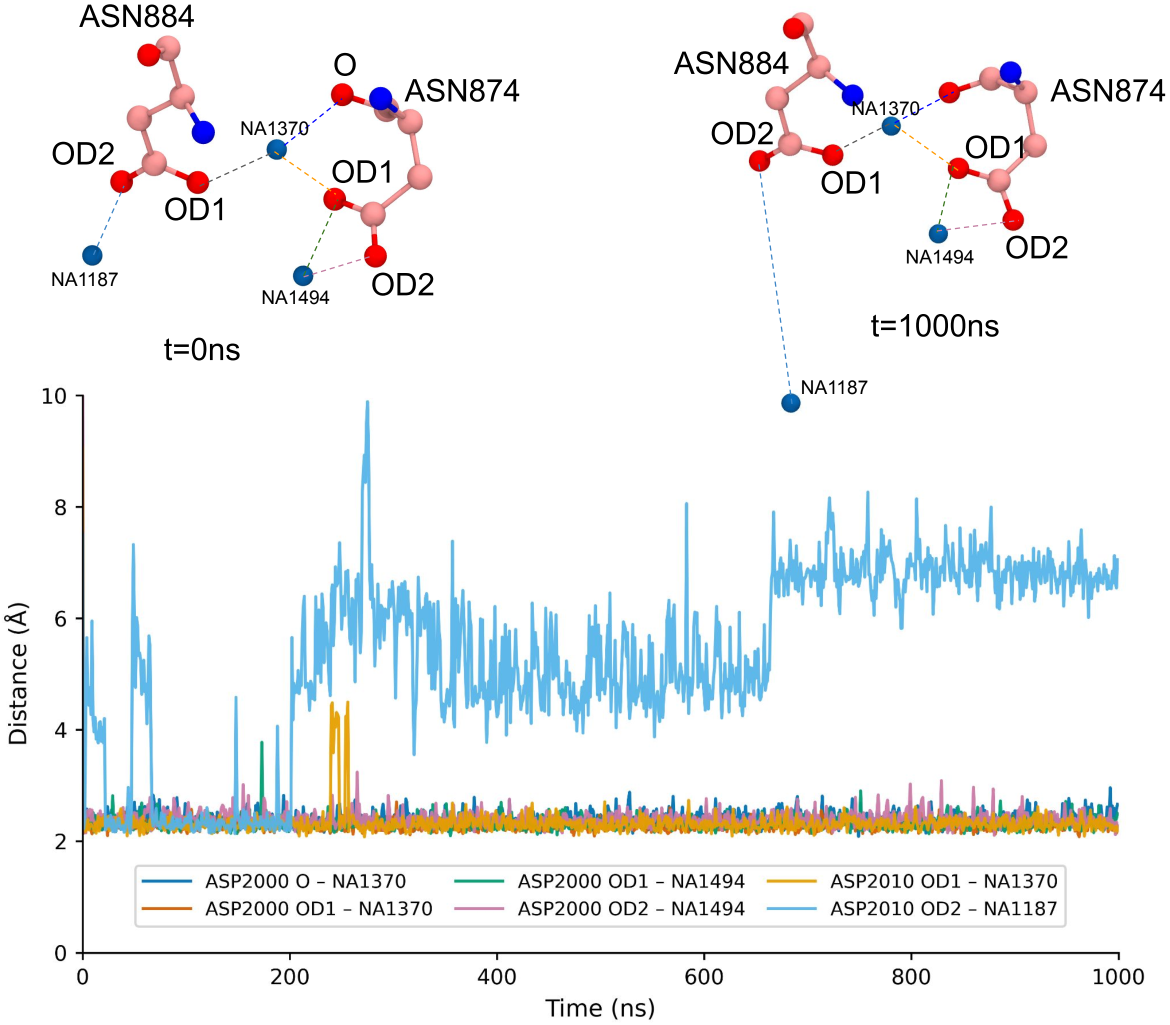


Figure S10: Visual representation and time series of multiple $Na^+$-mediated bridging event between two Asp residues, involving both side-chain carboxylate oxygen atoms and backbone oxygen atoms. Shaded areas indicate the interface between the dense and dilute phases.

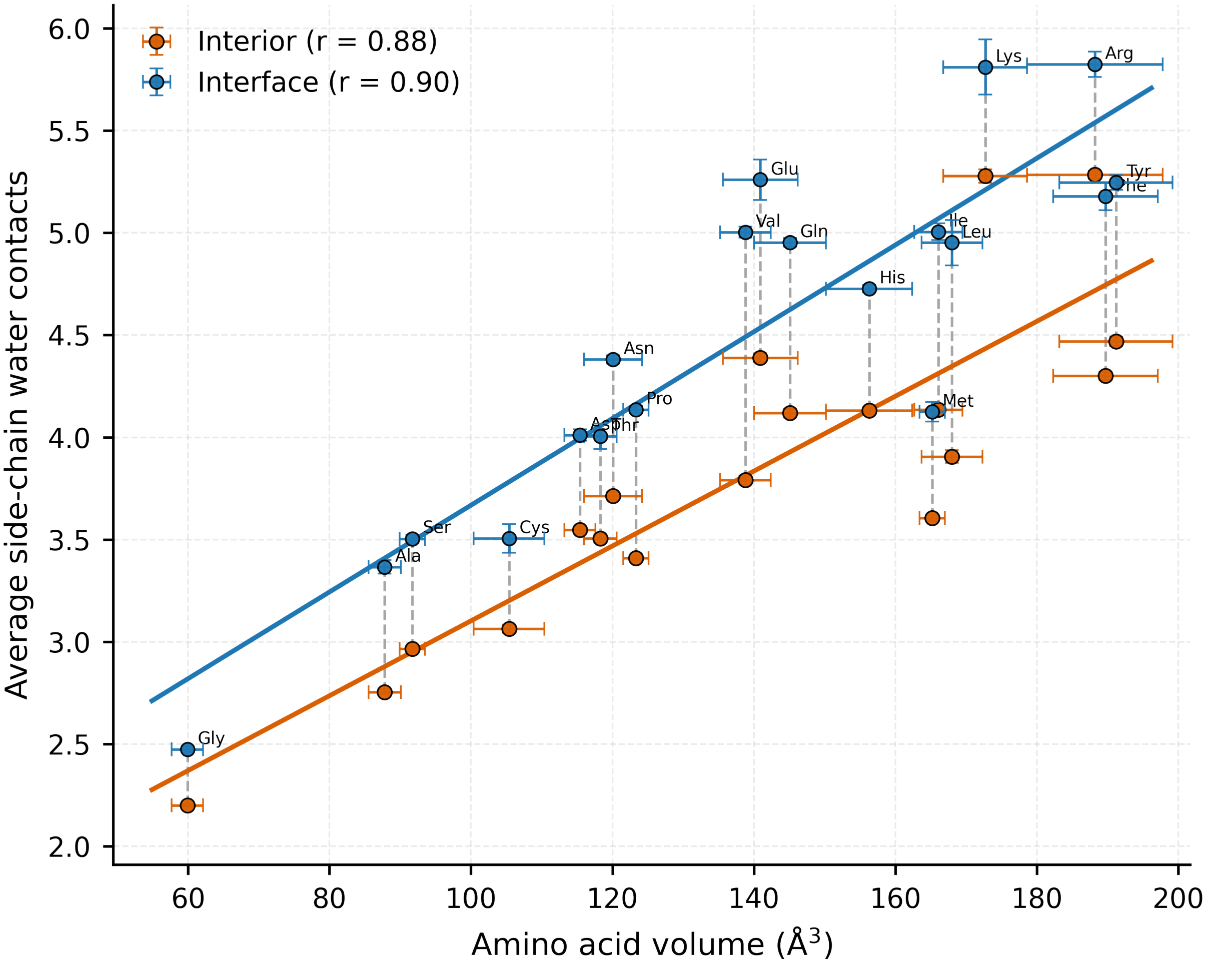


Figure S11: Correlation plot illustrating the relationship between the average number of water molecules interacting with amino acid side chains and the corresponding side-chain volume, evaluated separately for residues located in the condensate interior and in the bulk phase.
